# The guardian of the genome meets a viral master gene regulator at a biomolecular condensate

**DOI:** 10.1101/2022.02.09.479752

**Authors:** Silvia Susana Borkosky, Marisol Fassolari, Karen Campos-León, Andrés Hugo Rossi, Mariano Salgueiro, Carla Pascuale, Ramón Peralta Martínez, Kevin Gaston, Gonzalo de Prat Gay

**Affiliations:** Fundación Instituto Leloir, Instituto de Investigaciones Bioquímicas de Buenos Aires (IIB-BA)-CONICET, Av. Patricias Argentinas 435, (1405) Buenos Aires, Argentina; Fundación para Investigaciones Biológicas Aplicadas (FIBA, Instituto de Investigaciones en Biodiversidad y Biotecnología (INBIOTEC)-CONICET, Mar del Plata, Argentina; School of Medicine, University of Nottingham Biodiscovery Institute, Nottingham, United Kingdom; Division of Immunity and Infection, School of Medicine, University of Birmingham, United Kingdom

## Abstract

As guardian of the genome, p53 exerts its tumor suppressor activity by modulating the expression of several hundreds of genes and by interacting with a large number of proteins. However, p53 can also repress viral replication and it is targeted by a variety of viral proteins to allow viral replication to proceed. p53 can repress human papillomavirus replication by binding to the viral E2 master gene regulator. Here we show how full-length p53 can spontaneously form phase separated liquid-like droplets that evolve to amyloid-like aggregates in a time-dependent manner, highlighting the fact that homotypic condensation is on the path to aggregation as observed in several protein aggregopathies. The DNA binding domain of HPV E2 (E2C) triggers heterotypic liquid-liquid phase separation with p53 with a precise 1 p53 : 2 E2C stoichiometry at the onset for demixing, yielding large regular spherical droplets that increase in size with E2C concentration. Moreover, E2C is able to slowly reshape time-evolved p53 aggregates into regular heterotypic liquid droplets. Using *in situ* sub-cellular fractionation, we show that E2 and wild-type p53 co-localize to the nucleus with a grainy pattern, and E2 can re-localize p53 into chromatin associated foci, a function independent of the DNA binding capacity of p53. A small DNA duplex containing the specific binding site for p53 deforms and dissolves both homotypic and heterotypic condensates at a 1 p53 : 1 DNA stoichiometry, whereas a ∼1000 base pair DNA fragment instead reshaped the condensates into distinct amorphous condensates containing p53, E2C and DNA, reminiscent of what we observe bound to chromatin. We conclude that p53 is a scaffold for liquid-liquid phase separation in line with its structural and functional features, in particular as a hub that binds multiple cellular protein partners as well as nucleic acids. Moreover, the capacity of E2C to rescue p53 from the amyloid aggregation route impacts on p53-rescuing drugs cancers where p53 mutation leads to loss of function.

## INTRODUCTION

Tumor suppressor p53 protein is a global transcription factor important for the prevention of cell transformation and ultimately cancer progression. In response to oncogenic or other types of cellular stress, p53 binds to specific DNA sequences to trigger up- or down-regulation of a variety of genes involved in cell cycle arrest, DNA repair, senescence, or apoptosis (Levine et al., 2006; Vogelstein et al., 2000). A variety of stress signals are known to induce p53 activity, including DNA damage, hypoxia, oncogene activation, inflammation, metabolic stress, and viral infection (Blagih et al., 2020; Labuschagne et al., 2018; Levine et al., 2006). The structural organization of p53 reflects the protein functional complexity, i.e., a modular domain structure, consisting of independently folded DNA-binding and tetramerization domains and intrinsically disordered regions that account for about 40% of the full-length protein (Joerger and Fersht, 2008). These disordered regions include the negatively charged N-terminal transactivation domain (p53_TAD_), a short proline rich region, and the C-terminal regulatory domain (p53_CT_) (Joerger and Fersht, 2008). In around 60% of human cancers, p53 is directly inactivated by mutation, with most mutational hot spots located within the DNA binding, also referred to as the “core” domain (p53_DBD_) (Joerger and Fersht, 2007, 2016). These hot spot mutations either decrease p53 thermodynamic stability or affect DNA-binding activity depending on their location (Joerger and Fersht, 2007). Mutations can impact on the functionality of p53 either leading to a loss of function (LOF) phenotype, a dominant negative (DN) effect on the remaining wild-type p53, or even a gain of function (GOF) activity (Bargonetti and Prives, 2019; Pilley et al., 2021; Sabapathy and Lane, 2018). Aside of the instability of the mutants, wild-type p53 aggregates can form amyloid-like homo and hetero aggregates with mutant p53 (Joerger and Fersht, 2016) with direct impact on p53 activity (de Oliveira et al., 2020). The p53_DBD_ denaturation takes place at near physiological temperature and governs the overall stability of the full-length protein since it ultimately leads to irreversible aggregation (Bullock et al., 1997) giving rise to different morphologies, including amorphous, fibrillar, and prion-like, depending on the experimental conditions (Ano Bom et al., 2012; Ishimaru et al., 2003). Therefore, based on the central role of p53 mutant conformationally instability and consequent aggregation on cancer development, several small molecule and peptide inhibitors of p53 aggregation designed to restore p53 native conformation have been explored as promising therapeutic strategies (Boeckler et al., 2008; Soragni et al., 2016).

Liquid–liquid phase separation (LLPS) or biomolecular condensation underlies the formation of membrane-less organelles providing a mechanism to dynamically compartmentalize the intracellular space (Banani et al., 2017; Hyman et al., 2014; Shin and Brangwynne, 2017). Hydrophobic and electrostatic interactions driven by intrinsically disordered regions (IDRs) in proteins, weak multivalent interactions, and nucleic acid binding all promote LLPS (Kaur et al., 2021; Li et al., 2012). A multimutated p53 variant was shown to undergo homotypic LLPS, and this was proposed to regulate p53 function (Kamagata et al., 2020). Condensates composed of proteins rich in IDRs are known to mature from liquid-like to solid-like sates, particularly those related to aggregopathies (Alberti and Dormann, 2019; Banani et al., 2017). A recent study showed that under strong crowding conditions the p53_DBD_ phase-separates *in vitro* and progresses into amyloid-like aggregation and described liquid-like GFP-p53 condensates in the nucleus (Petronilho et al., 2021). Another report described anomalous and amorphous homotypic condensates of p53 hosting anomalous amyloid fibrillar aggregates (Safari et al., 2019). Although p53 LLPS was proposed to be a precursor or intermediate state on the amyloid aggregation pathway, direct sequential mechanistic evidence is still lacking.

Human papillomaviruses (HPVs) infect epithelial cells and generally induce the formation of benign hyperproliferative lesions. However, a subset of these HPVs is known to be oncogenic, and persistent infection with these high-risk HPVs can result in anogenital or oropharyngeal cancers (Schiffman et al., 2016). The HPV E2 protein is the master regulator of the virus life cycle, with main roles involving the control of gene transcription and viral genome replication (McBride, 2013). E2 role in transcription is well characterized, it functions primarily by recruiting cellular factors to the viral genomes, which activate or repress transcriptional processes upon specific binding to the viral genome in a dose-dependent manner (McBride, 2013; Warburton et al., 2021). Additionally, E2 acts as a loading factor for the viral helicase E1 at the HPV origin of replication and in late stages of replication has crucial roles in genome maintenance and partition (Warburton et al., 2021). The E2 proteins are around 400 amino acid polypeptides consisting of an N-terminal transactivation domain (ca. 200 amino acids) and a C-terminal DNA binding and dimerization domain (E2C, ca. 85 amino acids), separated by a non-conserved “hinge” domain. The E2C domain folds as an obligated dimeric β-barrel, consisting of 2 half-barrels each composed of 4 β-strands and 4 α-helices: one α-helix from each half-barrel acts as a DNA recognition site (de Prat-Gay et al., 2008; Hegde, 2002).

p53 discovery was linked to early investigations on how DNA tumor viruses cause cancer, and demonstrate that proteins from three distinct tumor viruses – SV40 T-antigen, E1B-58K adenovirus protein, and the HPV E6 oncoprotein– formed complexes with p53 (Vogelstein et al., 2000). Interestingly, p53 was subsequently found at the viral replication sites of SV40, herpes simplex virus (HSV), cytomegalovirus (CMV) and adenovirus (Fortunato and Spector, 1998; Gannon and Lane, 1987; Konig et al., 1999; Wilcock and Lane, 1991). Replication of HPV is carried out at so-called replication foci or HPV E1/E2 foci, containing the viral helicase E1 and E2 (Warburton et al., 2021). These E1/E2 foci represent viral replication factories, and they recruit DNA damage response (DDR) proteins including p53 phosphorylated at serine 15 (Sakakibara et al., 2011; Warburton et al., 2021). E2 protein was shown to bind to p53 by pull-down assays *in vitro* and in cells (Massimi et al., 1999; Parish et al., 2006). Moreover, p53 has been shown to target E2 to inhibit HPV16 DNA replication and transcriptional activity (Brown et al., 2008).

In this work we show that full-length pseudo wild-type p53 spontaneously forms homotypic liquid-like droplets on-pathway to amyloid-like aggregates. We find that the interaction with HPV E2 that takes place in cells is based on heterotypic phase separation, and that the viral master regulator not only prevents but readily rescues the p53 aggregates to yield highly regular liquid droplets. While a small DNA duplex with a p53 binding site dissolves the droplets, long DNA fragments reshapes the latter into aggregates containing both proteins and the DNA, consistent with the co-localization of both proteins at chromatin associated foci that we here describe. We show that p53 acts as an LLPS scaffolder and propose this feature as the basis for its role as a cellular hub arising from its well documented binding promiscuity. This work uncovers two biologically meaningful events behind p53 LLPS: homotypic LLPS as intermediate in the cancer aggregation route resembling neuronal aggregopathies, and heterotypic LLPS linked to a known functional interplay with the E2 master regulator. We discuss the biological implications and impact on p53 rescuing therapeutics.

## RESULTS

### Formation of p53 ionic strength-dependent droplets is enhanced by molecular crowding

Wild-type full-length p53 is known for a marked tendency to readily undergo thermal aggregation in mild physiological conditions (t_m_ 42 °C) (Nikolova et al., 1998). In order to address the capacity of p53 to form condensates, we used a rationally designed stability mutant of the full-length protein, previously described as pseudo wild-type p53 (Nikolova et al., 1998). Transfer of fluorescently labeled cy5-p53 from ice to room temperature, pH 7.0, and low salt led to an increase in turbidity reaching a plateau at 30 minutes, which corresponded to the formation of small (1-2 μm diameter) regular droplets as judged by both fluorescence and bright field (BF) microscopy (Figure 1A). Subsequent addition of 200 mM NaCl completely reverted both the scattering signal and the formation of droplets (Figure 1A). To assess the effect of molecular crowding, a comparable experiment was performed with the addition of 5% PEG. This led to a more pronounced change in the absorbance scattering signal corresponding to the formation of larger and highly regular spherical droplets, of up to 7 μm in diameter (Figure 1B). Addition of 200 mM NaCl also dissolved the droplets, indicating that droplet condensation is governed by ionic strength in the presence and absence of crowding. Although increasing the temperature raised the rate of condensation, the same extent of signal was reached at either 10 or 45 °C, suggesting no effect on the steady state condensation (Figure S1A). However, the species formed at 10 °C completely reverted upon addition of salt while the recovery at 45 °C was only partial (Figure S1A), most likely due to irreversible aggregation, considering a t_m_ of 47.2 °C for the p53 pseudo wild-type protein (Nikolova et al., 1998).

**Figure 1.**
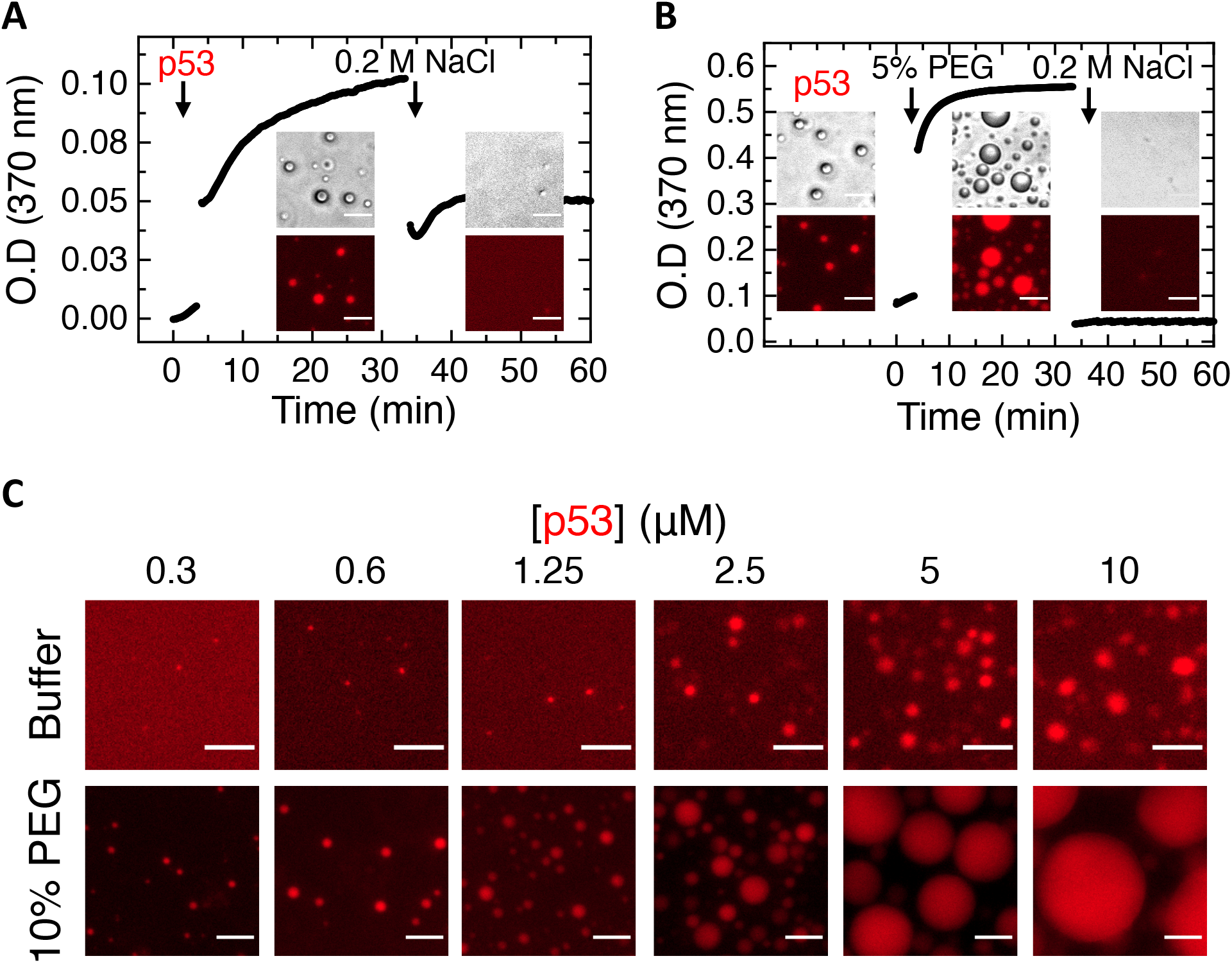
p53 forms spherical droplets. (A) Turbidity kinetic assay of samples containing cy5-labelled p53 (Cy5-p53) monitored by light scattering combined with visualization by bright field and fluorescence microscopy; NaCl was added after the signal reached maximum absorbance. (B) Similar experiment as in A but in presence of crowding agent (5% PEG). (C) Representative images of samples with different concentrations of cy5-p53 in absence or presence of PEG visualized by fluorescence microscopy. Scale bars= 10 μm.

Using fluorescence microscopy, we observed an increase in the size of the droplets depending on protein concentration over the range tested, reaching a maximum diameter of 3 μm and 30 μm in the absence or presence of crowding agent, respectively (Figure 1C). This concentration effect in the presence of PEG was clearly evidenced by turbidity assays (Figure S1B). The onset for the formation of the condensates under crowding conditions is ∼0.6 μM compared to ∼2.5 μM p53 tetramer concentration in the absence of crowding agent (Figures 1C). Inspection of the images suggests that besides a size limit without the crowding agent, the aspect of the droplets is different as evidenced by fluorescence microscopy (Figure 1C). In control experiments, we observed that droplet formation was independent of the fluorescently label (Figure S1C).

### Homotypic liquid-liquid phase separation droplets of p53 evolve to aggregates with amyloid properties

Having shown that p53 forms regular reversible droplets modulated by protein concentration and molecular crowding, we aimed at evaluating its material properties using fluorescence recovery after photobleaching (FRAP) experiments. After bleaching, 70 % of the total signal was recovered, with a major exponential phase governing the process, with a *t*_1/2_ of 80 seconds (Figures 2A, S2A and S2B). Nevertheless, the droplets displayed incomplete coalescence into non-regular clusters, suggesting some sort of maturation over longer periods (Figure S2C). To further analyze the maturation of the p53 homotypic droplets, we carried out time course experiments at room temperature. Regular spherical droplets observed in the absence of crowding agent within the first hour evolved to clusters of droplets, clearly different from amorphous aggregates (Figures 2B left, S2D and S2E). However, in the presence of PEG, the droplets remained highly spherical, increasing in size with time (Figures 2B right, S2D and S2E). Identical results were found when FITC was used instead of cy5 (Figure S2E). Since it has been described that p53 undergoes aggregation through the formation of amyloid-like intermediates (Ano Bom et al., 2012; Ishimaru et al., 2003), we considered the evolution of the initial spontaneous homotypic droplets into an aggregation route. For this, we carried out similar time course experiments, and added thioflavin T (ThT) as an indicator of amyloid-like repetitive β-sheet structure. Indeed, an increase in ThT fluorescence was observed in parallel to the formation of the droplets and coalescence into the same clusters with p53, a process completed at five hours (Figure 2B). In the presence of PEG, an homogeneously distributed ThT staining of the droplet suggests that conformational species with repetitive β-sheet conformations typical of amyloid-like structures are present (Figure 2B, right). This is supported by the fact that the crowding agent stabilized droplets showed no recovery in FRAP experiments (Not shown). Thus, p53 homotypic condensates are liquid in nature but show slow dynamics and evolution to maturation through a β-amyloid aggregation route. While the intensity of the protein cy5 fluorescence is similar in both processes, the much lower intensity of the ThT labelling in the presence of crowding agent suggests a different type of conformation in the PEG stabilized droplet (Figure 2B, right, 24 hrs).

**Figure 2.**
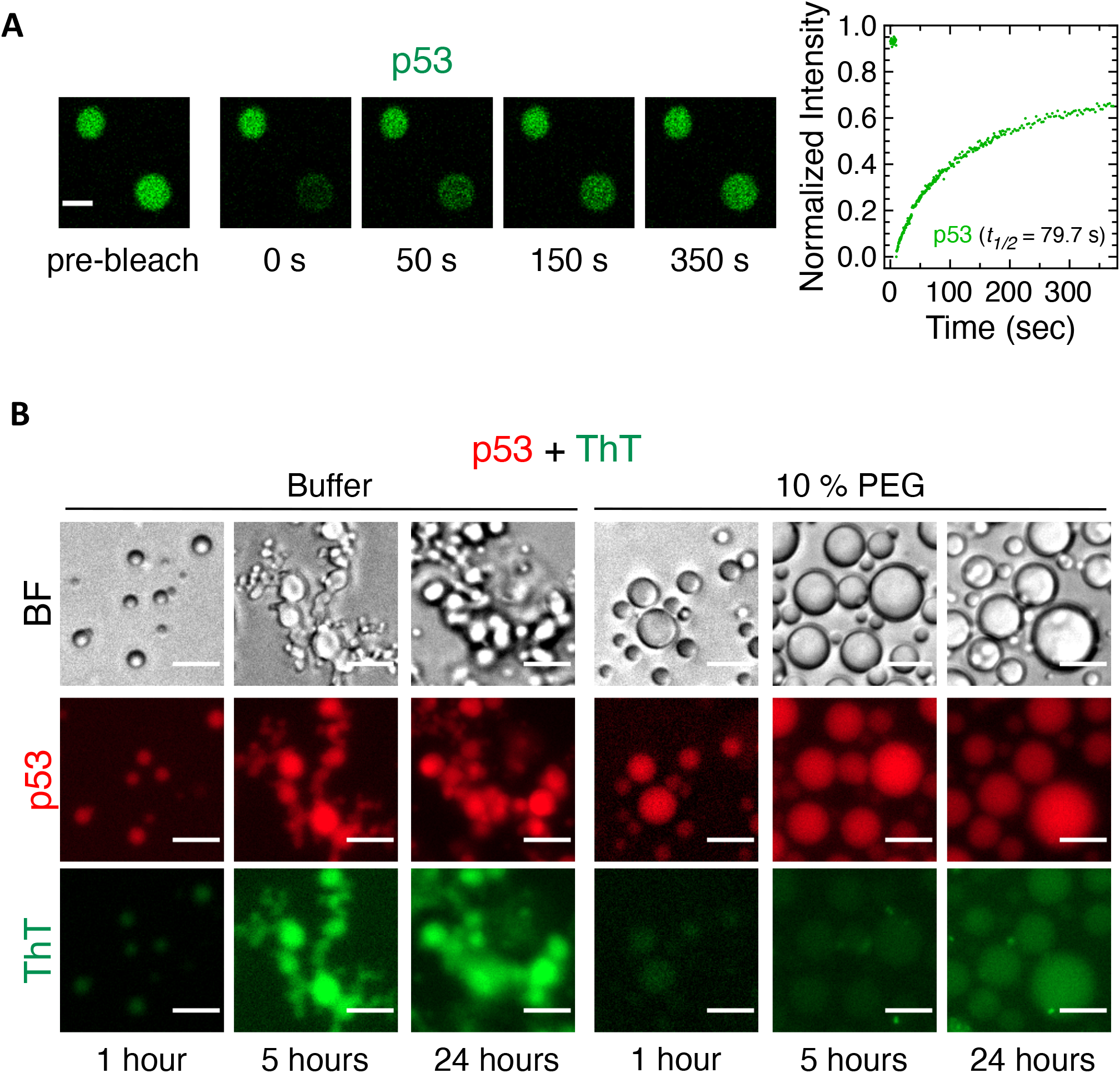
p53 droplets are liquid-like and evolve to amyloid-like structures. (A) Representative confocal microscopy images and mean plot of fluorescence recovery after photobleaching (FRAP) analysis of homotypic droplets composed of 2.5 μM FITC-labelled p53 (FITC-p53) (n= 7 droplets). Data were normalized to the average intensity of a droplet not photobleached and fitted using a double exponential function. Half time recovery (*t*_*1/2*_) of 79.7 ± 11.5 seconds was obtained. Scale bars= 2 μm. (B) Representative microscopy images of samples containing cy5-p53 incubated with Thioflavin T (ThT) during 1, 5 and 24 hours, without crowding agent (Left panel) or in presence of 10% PEG (Right panel). BF, bright field. Scale bars= 10 μm.

### Heterotypic condensation of p53 with HPV16 E2C is finely tuned by stoichiometry

Given the functional interplay between p53 and HPV E2 (Brown et al., 2008; Desaintes et al., 1999; Massimi et al., 1999; Parish et al., 2006; Webster et al., 2000), we addressed the nature of the interaction between the two proteins *in vitro*. The interaction with p53 was mapped to the C-terminal DNA-binding domain of the HPV16 E2 protein (E2C) (Massimi et al., 1999; Parish et al., 2006). It should be noted that p53 is a tetramer with *K*_*D*_ of 50 nM (Rajagopalan et al., 2011) while E2C is an obligated dimer with *K*_*D*_ of 30 nM (Mok et al., 1996), thus, we refer to tetramer concentration for p53 and dimer concentration of E2C throughout this work. When both proteins were co-incubated at different concentration ratios using a fixed p53 concentration of 0.25 μM, we observed a kinetic increase in absorbance scattering at 370 nm as a function of E2C concentration (Figure 3A and B), and SDS-PAGE analysis of soluble and insoluble fractions showed a stoichiometric formation of a centrifugable material at a 2:1 to 4:1 E2C:p53 ratio (Figure 3B).

**Figure 3.**
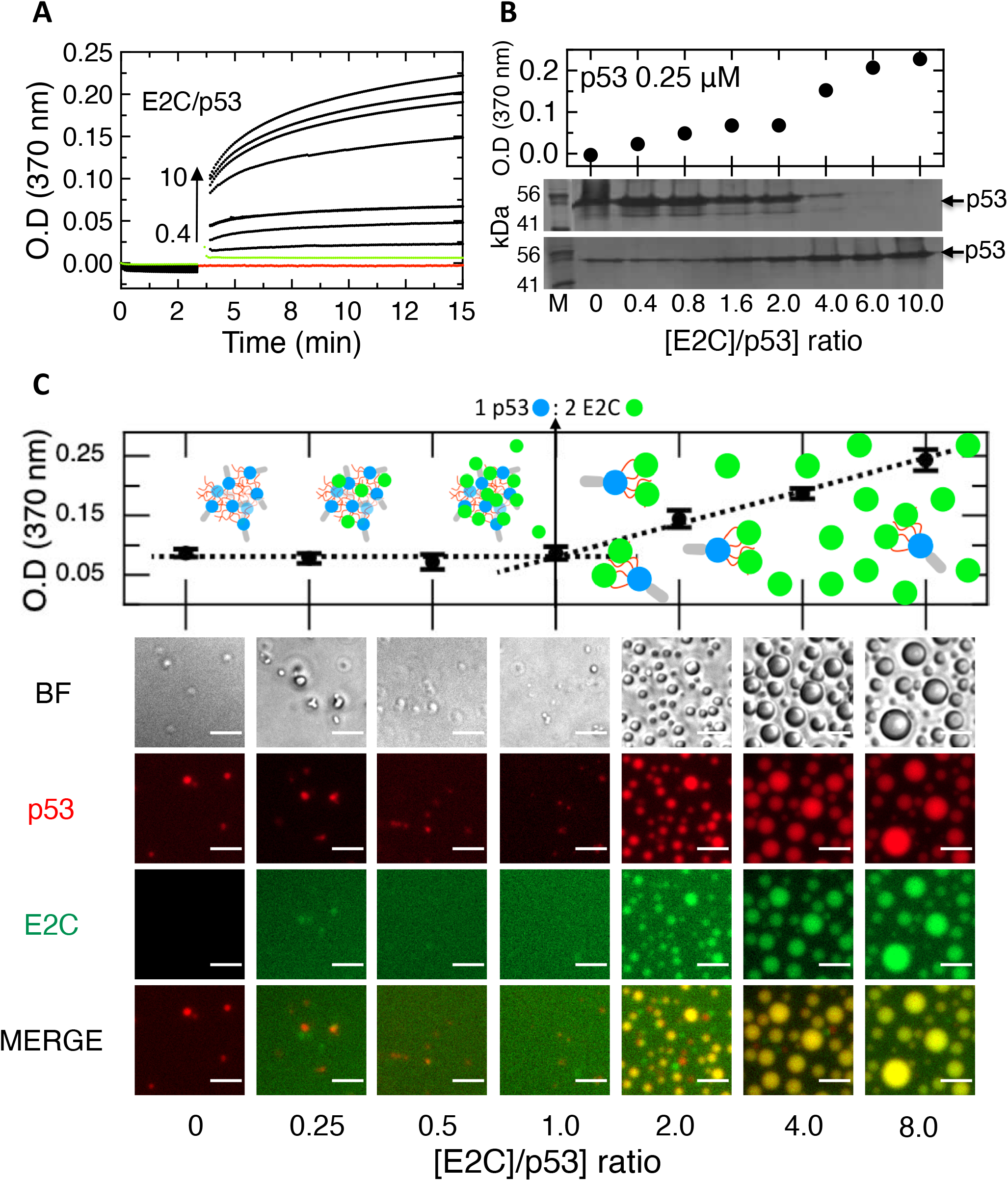
p53 interaction with HPV16 E2C leads to heterotypic regular droplets. (A) Kinetic light scattering plot of individual samples containing 0.25 μM p53 incubated for 3 minutes, followed by addition of increasing concentrations of HPV16 E2C (Black dots). Samples with E2C/p53 ratios of 0.4, 0.8, 1.6, 2, 4, 6 and 10 were included. Control samples contained 0.25 μM p53 (Red dots) or 2.5 μM E2C (Green dots). (B) Plot of the final points of the kinetic assay from A (Upper panel). Soluble (Upper gel image) and insoluble (Lower gel image) fractions of the samples obtained by centrifugation and ran by SDS-PAGE electrophoresis (Lower panel). M, molecular weight marker. (C) Light scattering assay monitoring absorbance at 370 nm on individual samples containing 2.5 μM cy5-p53 and increasing FITC-labelled E2C (FITC-E2C) concentrations. Blue circles correspond to p53 DBD, red chains to p53 TAD, grey cylinders to p53 tet domain, and green circles to E2C (Upper panel). Representative microcopy images of samples measured in B and visualized after incubation for 1 hour (Lower panel). Yellow regions show colocalization in the merged images. BF, bright field. Scale bars= 10 μm.

The formation of scattering insoluble species upon co-incubation with E2C prompted us to consider a heterotypic LLPS process involving both proteins. We incubated fixed amounts of p53 (2.5 μM) with increasing concentrations of E2C. After 1 hour, a sharp onset of droplet formation at a 2:1 E2C:p53 ratio with complete co-localization of both proteins was observed, coincident with turbidity increment (Figure 3C). After a seven-hour incubation period, the samples with sub-stoichiometric amounts of E2C (0.25:1 to 1:1 E2C:p53 ratios) evolved to irregular bead-like structures, similar to the aggregates of p53 alone, with colocalization of both proteins (Figure S3A). From these results, we can conclude that stoichiometric amounts of E2C prevent the spontaneous aggregation route of p53. Control experiments showed that concentrated samples of E2C have not formed either aggregates or homotypic LLPS droplets in the absence of p53 (Figure S3B). In support for the liquid nature of the p53:E2C droplets, we observed coalescence events of the droplets relaxing to a spherical shape in less than 10 seconds (Figure S3C). Based on the observed stoichiometry, further experiments were performed using a fixed a concentration ratio of 4:1 E2C:p53 to ensure optimal conditions for heterotypic LLPS.

To evaluate the material properties of the heterotypic p53:E2C droplets we carried out FRAP analysis using FITC labelling on both proteins in separate individual experiments for a more consistent comparison (Figures 4A and S4A). p53 recovered 73% of the pre-bleach intensity, with a *t*_1/2_ of 63 seconds (Figure 4A), slightly faster than the recovery of p53 in homotypic droplets. E2C recovered to a similar extent (84%) but with a significantly faster recovery rate than that p53 (*t*_1/2_ 31 seconds) (Figure 4A).

**Figure 4.**
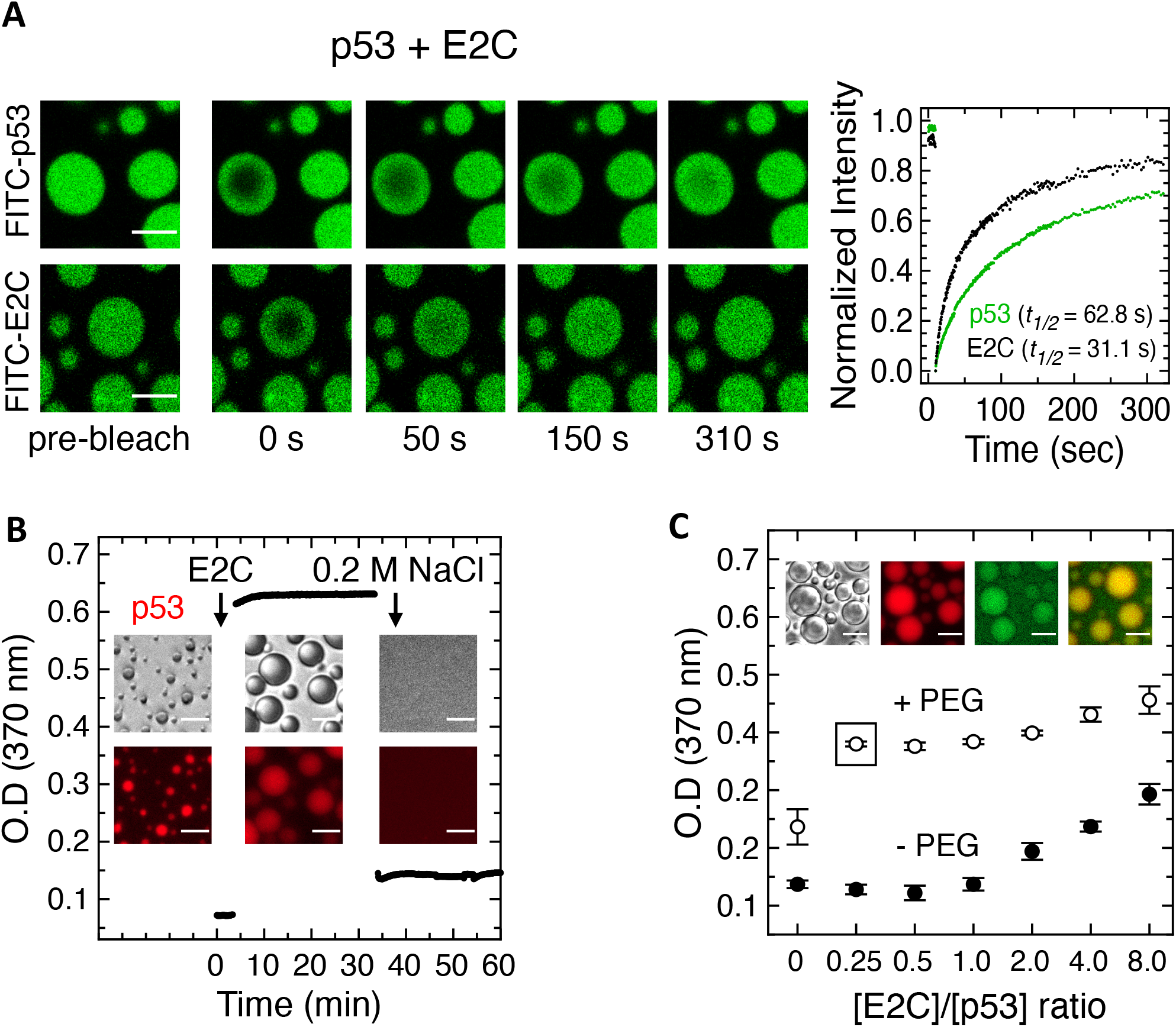
Material properties of p53:E2C heterotypic LLPS. (A) Representative confocal microscopy images and mean plot of FRAP analysis of heterotypic spherical droplets composed of 2.5 μM FITC-p53 and 10 μM unlabelled E2C (Upper image panel and green dots on plot) or 2.5 μM unlabelled p53 and 10 μM FITC-E2C (Lower image panel and black dots on plot). Data were normalized to the average intensity of a droplet not photobleached and fitted using a double exponential function (n (p53)= 7 droplets; n (E2C)= 8 droplets). Half time recoveries (*t*_*1/2*_) of 62.8 ± 10.8 seconds and 31.3 ± 3.3 seconds were obtained for p53 and E2C, respectively. Scale bars= 5 μm. (B) Cy5-p53 homotypic droplets screened by turbidity kinetic assay and visualized by BF and fluorescence microscopy. Heterotypic condensation was triggered by addition of E2C and reversed by increasing the ionic strength. (C) Light scattering assay monitoring absorbance at 370 nm of samples containing 2.5 μM cy5-p53 and increasing E2C concentrations with 10% PEG (open circles) in comparison to similar samples without crowding agent (Black circles) (Same samples as in figure 3C). Representative microscopy images of the samples with E2C/p53 ratio of 0.25 in presence of 10% PEG after incubation for 7 hours (Highlighted with box on plot). BF, bright field. Scale bars= 10 μm.

As p53 homotypic LLPS proved to be highly sensitive to salt, we sought to evaluate the effect of ionic strength on the heterotypic p53:E2C LLPS. We monitored the turbidity of a 2:1 mixture of E2C:p53 with increasing concentrations of NaCl, observing signal disappearance at 100 mM salt concentration (Figure S4B, left) with concomitant increase of protein soluble fraction – monitored by absorbance at 280 nm (Figure S4B, right). We confirmed the complete reversibility of this ionic strength-dependent process by performing a sequential kinetic assay, registering microscopy images thus validating the turbidity signal as LLPS. For this, E2C was added to the pre-formed typically small p53 homotypic droplets at low salt, giving rise to a large increase in absorbance and the formation of large spherical heterotypic droplets within the experimental deadtime (ca. 20 seconds); subsequent addition of 200 mM NaCl completely reversed both the LLPS droplets and turbidity (Figure 4B). Next, we evaluated the effect of molecular crowding on heterotypic p53:E2C LLPS. The onset for droplet formation in the presence of crowding agent dropped from 2:1 to 1:4 E2C:p53 ratio (Figure 4C). This is in line with the formation of large p53 homotypic droplets in the presence of crowding agent (Figure 1B and C), indicating minimal concentrations of E2C to be recruited into the droplets, as shown by protein co-localization (Figure 4C).

### E2C rescues p53 from the amyloid aggregation route

Having shown that p53 homotypic LLPS droplets gradually evolve into an amyloid-like aggregation route, we decided to address the effects of E2C on this process. To this end, p53 was incubated for 2 hours to allow for the formation of the bead-like clustered aggregates described above (Figure 5A). Following addition of E2C to the pre-incubated p53 sample at a 4:1 E2C:p53 ratio, the clusters were reshaped into highly regular spherical droplets which increased in size as time elapsed (Figure 5A, central panel). Control experiments without the addition of E2C at the same time points showed the presence of bead-like clustered as opposed to amorphous aggregates (Figure 5A, right panel). In addition, experiments combining unlabelled proteins, cy5-p53 with unlabeled E2C, or unlabeled p53 with FITC-E2C showed that the process is not affected by the presence of either fluorophore (Figure S5A, B and C). In a separate experiment, we reversed the order of mixture of the proteins, adding p53 without pre-incubation into a solution containing pre-incubated E2C. We observed that homotypic p53 droplets are formed within the first 5 minutes, indicating that the immediate formation of these droplets (Experimental dead time ca. 20 seconds, Figure 4B) is not affected by the presence of E2C in the solution (Figure S5D). Incorporation of E2C to the droplets was barely detectable at 20 min and complete at 3 hours (Figure S5D).

**Figure 5.**
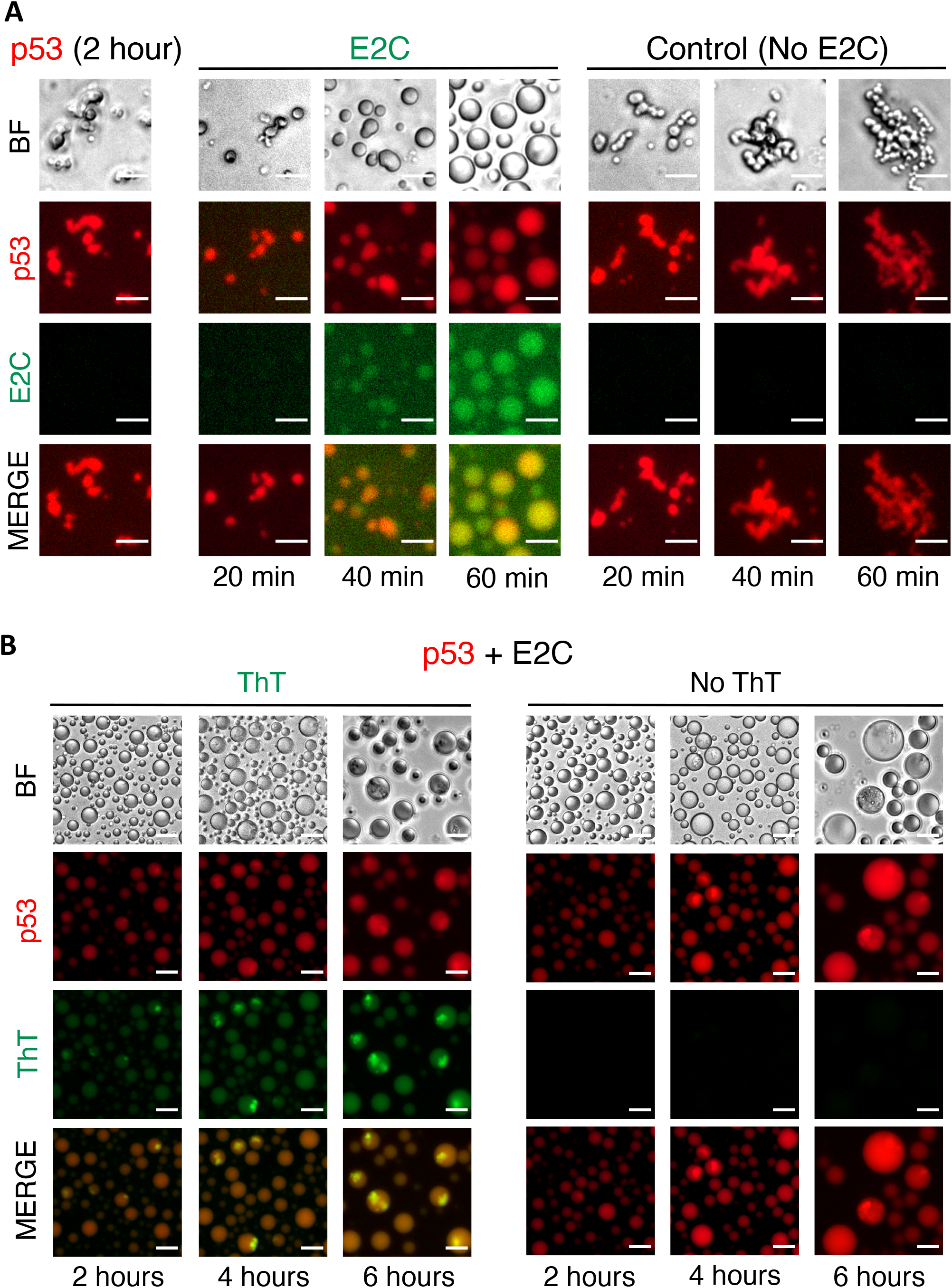
p53 aggregation fate is modulated by E2C. A) Representative microscopy images of a sample containing 2-hour old cy5-p53 irregularly shaped clusters (Left panel) followed by addition of FITC-E2C (Central panel). Time-course examination of the evolution of the clusters into spherical droplets at 20, 40 and 60 min (Central panel). Control cy5-p53 homotypic samples without addition of E2C (Right panel). (B) BF and fluorescence microscopy images of heterotypic droplets composed of cy5-p53 and unlabelled E2C incubated with ThT during 2, 4 and 6 hours (Left panel). Control samples without ThT (Right panel). BF, bright field. Scale bars= 20 μm.

We have shown that p53 spontaneously evolves into β-sheet amyloid-like aggregates (Figure 2B), and that E2C prevents and rescues the bead-like p53 aggregates by reshaping them into highly regular LLPS droplets (Figure 5A). To gain insight into the amyloid-like transitions within the rescued heterotypic droplets, we co-incubated a p53-E2C mixture with ThT and monitored the samples by microscopy at different incubation times. At two hours we observed diffuse homogeneous staining of ThT in all droplets, matching the p53 homogeneous distribution within the heterotypic droplet (Figure 5B). Interestingly, as time progressed a strong signal was detected for ThT-bound p53 aggregates partitioned within the droplets. These aggregates remained within the droplet boundaries, indicating that their formation occurred within the dense phase. The number of droplets containing these partitioned aggregates increased with time, suggesting that despite a prolonged delay of the aggregation route by E2C, the overall material properties of the heterotypic droplets caused the accumulation of β-sheet-rich aggregates inside the droplets. The formation of sub-compartments within liquid BMCs and viral replication complexes (VRCs) has been reported (Banani et al., 2017; Charman and Weitzman, 2020). As ThT binding structures partitioned within heterotypic droplets and did not evolve to large aggregates as observed for homotypic condensates (Figure 2B), we believe that within ageing heterotypic droplets, at least two compartmentalized populations exist: i) amyloid-like p53 aggregates; and ii) phase-separated complexes of p53 and E2C with dynamic behaviour.

### HPV16 E2 recruits p53 to chromatin foci

To evaluate whether condensed structures of p53 and HPV16 E2 form in cells we made use of transfection experiments. We first addressed colocalization by expressing wild-type p53 and HPV16 GFP-tagged E2 fusion protein in cells that lack endogenous p53 (p53-null Saos-2 cells). In transiently transfected Saos-2 cells, the subcellular localization of p53 is clearly nuclear while GFP-E2 is predominantly nuclear with some cytoplasmic signal (Figure 6A). Co-expression revealed that the GFP-E2 protein completely colocalized with p53 in the nuclei of these cells with a homogenously dense pattern (Figure 6A). We next wanted to address the nature of structures formed by interaction of the two proteins within the nucleus. For this, we used C-33A cervical keratinocytes, a cell line that expresses the p53 R273C mutant, widely documented for being impaired for DNA binding (Joerger and Fersht, 2007). This allowed us to examine an interaction between E2 and p53 in the absence of p53-DNA interaction, a determinant for its main functional and primary role in transcriptional activity. C-33A cells were transiently transfected with GFP-E2, and both proteins were monitored by immunofluorescence. Endogenous p53 R273C showed a fine grainy staining pattern homogeneously distributed in the nucleus (Figure 6B, left). A similar grainy pattern was observed when cells were transfected with GFP-E2, and the merge image shows that they indeed fully colocalized (Figure 6B, left). To further examine this localization, we performed *in situ* sub-cellular fractionation (Sawasdichai et al., 2010), where cytoplasmic and loosely held nuclear proteins are removed and only tightly chromatin bound proteins remain. When C-33A cells expressing GFP-E2 were subjected to *in situ* sub-cellular fractionation, E2 and p53 colocalized to chromatin-associated condensate-like material with a coarse granular pattern (Fig. 6B, right). However, in the absence of E2, p53 R273C was not retained in the chromatin fraction (Figure 6B, right), i.e., it is not tightly bound to chromatin.

**Figure 6.**
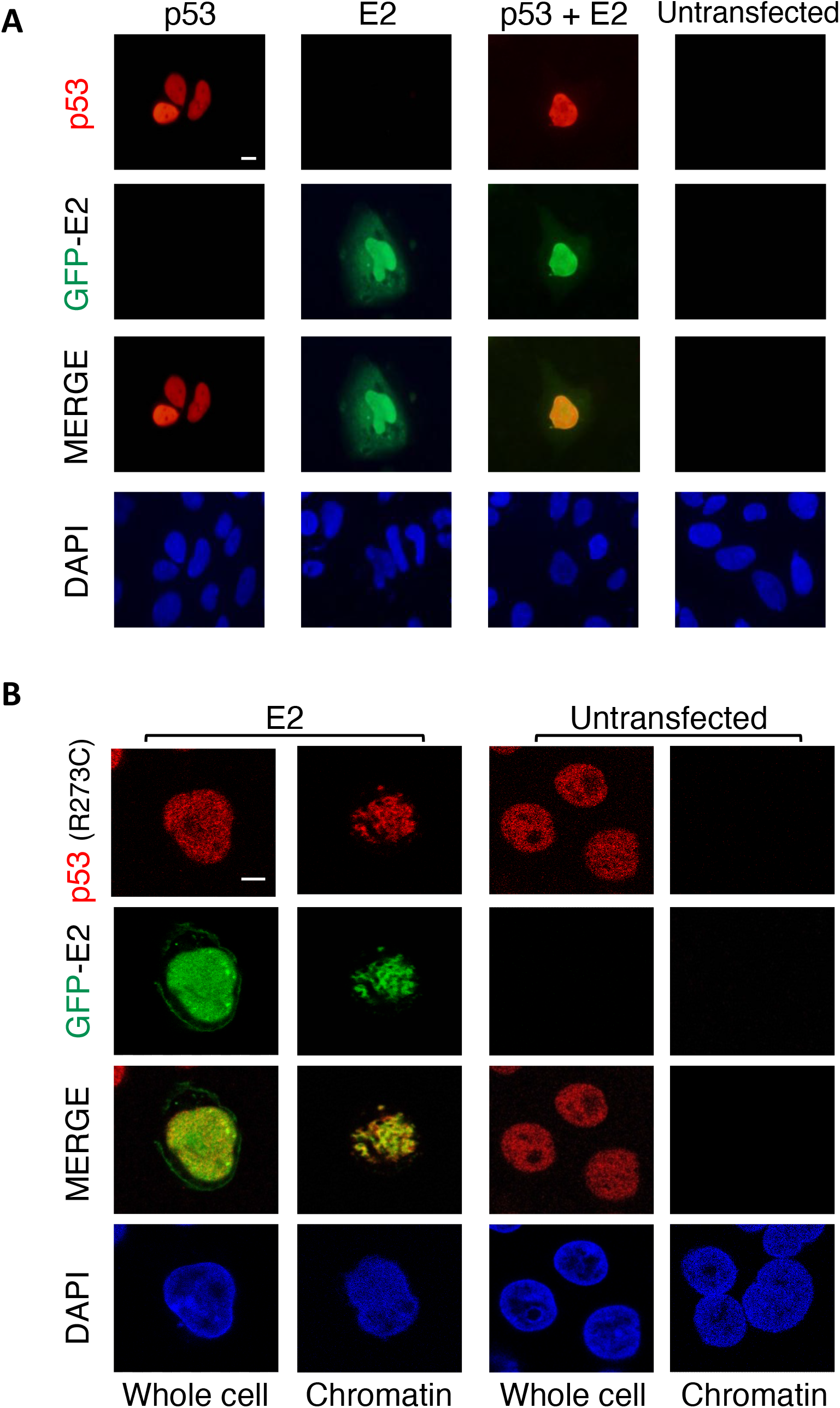
Co-localization of p53 and E2 in cells. (A) Fluorescence microscopy images of Saos-2 cells transfected with pCB6:p53 alone or in combination with pGFP HPV16 E2; protein expression was detected by immunofluoresence. GFP-E2 fusion protein and p53 were visualized with FITC and TRITC filters, respectively. (B) Confocal microscopy images of C-33A cells transiently transfected with pGFP E2 followed by immunofluorescence. Additionally, replicates of transfected and untransfected controls were subjected to *in situ* sub-cellular fractionation prior to immunofluorescence detection of E2 and p53 R273C, allowing for comparison between protein sub-cellular distribution within whole cells and in the chromatin fraction. GFP-E2 fusion protein and p53 were visualized using FITC and TRITC filters, respectively. Co-localization is demonstrated by yellow/light green regions in the merged images. DAPI staining was used to visualize the cell nuclei. Scale bars= 5 μm.

### Differential effect of a short p53 specific DNA duplex and a long non-specific DNA fragment on dissolution and reshaping of homotypic and heterotypic p53 condensates

The fact that both p53 and E2 are transcriptional regulators that interact and colocalize to chromatin condensate-like foci, prompted us to investigate the effect of DNA on the homotypic and heterotypic condensation processes *in vitro*. First, we incubated p53 with increasing concentrations of a 20 bp DNA duplex containing a cognate high affinity p53 binding site (el-Deiry et al., 1992), that we named DNA_p53_. Sub-stoichiometric amounts of DNA_p53_ (0.25:1 DNA:p53) produced deformation of the droplets which were ultimately completely dissolved at a 1:1 ratio (Figures 7A and S6A). These results indicate that the homotypic condensation of p53 relies on a self-interaction that involves its DNA binding site. Alternatively, DNA might induce conformational changes that affect self-interaction. Interestingly, when DNA_p53_ was added to the heterotypic p53-E2C droplets, we observed that sub-stoichiometric amounts of DNA gradually deformed and converted the regular droplets into dense mixed bead-like and amorphous aggregates that contained both proteins (Figures 7B, S6B and S6C). These aggregates first dispersed at intermediate DNA:p53 ratios, and completely dissolved at a 4:1 DNA:p53 ratio (Figures 7B and S6C), four-fold higher than the 1:1 ratio required for dissolution of homotypic p53 droplets (Figures 7A and S6A). Such an excess of DNA required for condensate dissolution strongly suggests that stoichiometric binding of DNA_p53_ to excess E2C (4:1 E2C:p53 for optimal heterotypic droplet formation) is linked to the fact that interactions made by the

**Figure 7.**
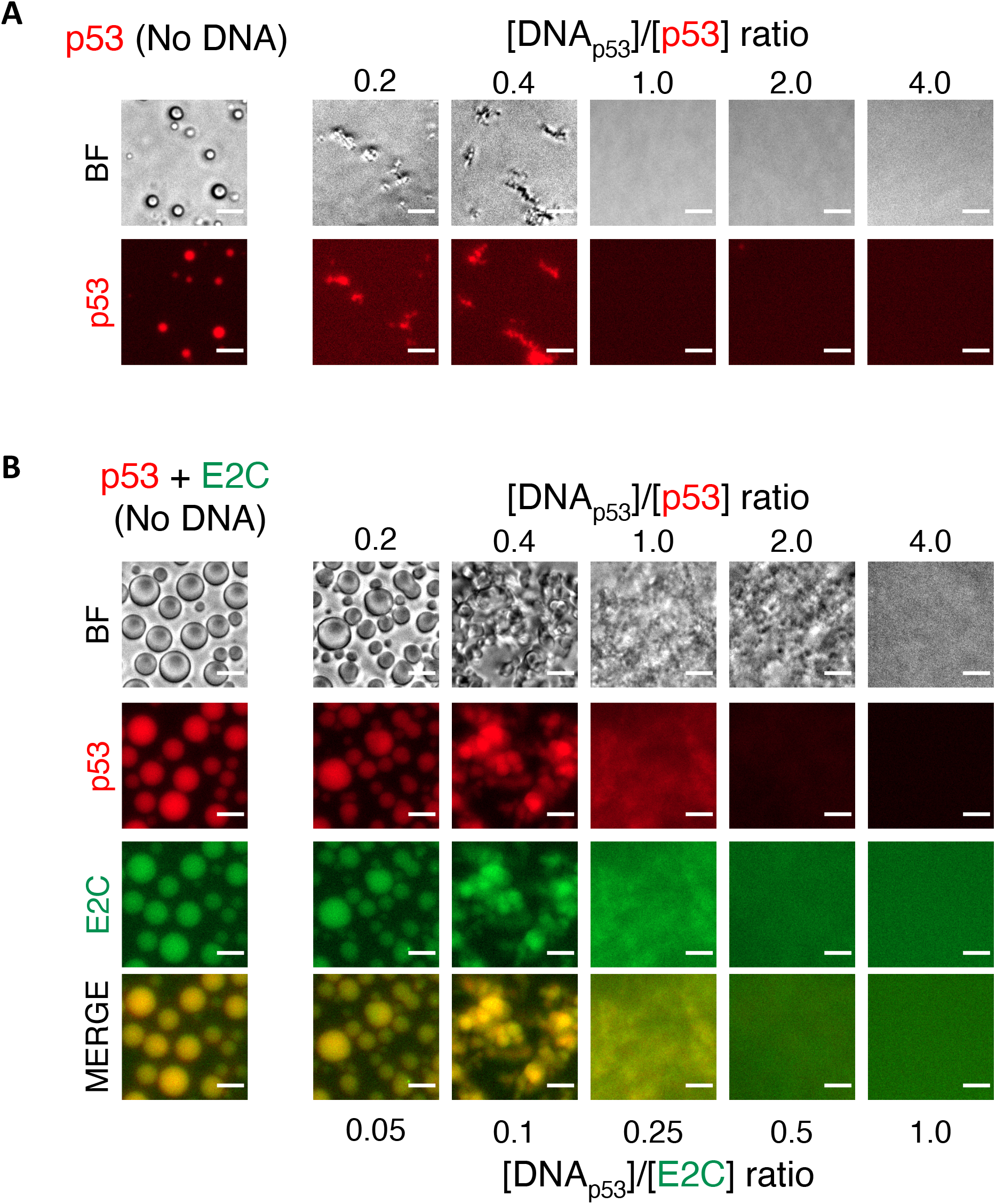
Effects of short oligonucleotide carrying a specific p53 binding site on LLPS. (A) Representative microscopy images of samples containing increasing concentrations of 20 bp dsDNA for p53 (DNA_p53_) and 2.5 μM cy5-p53. Control sample without DNA contained 2.5 μM cy5-p53. (B) Microscopy images of samples containing increasing concentrations of DNA_p53_, and fixed concentration of cy5-p53 and FITC-E2C (E2C/p53 ratio=4.0). Control sample without DNA contained 2.5 μM cy5-p53 and 10 μM FITC-E2C. Yellow regions show colocalization of p53 and E2C in the merged images. BF, bright field. Scale bars= 10 μm.

DNA binding site of E2C are involved in stabilizing the heterotypic p53:E2C droplets. The *K*_D_ for E2C binding to non-specific DNA is ca. 1 μM, compared to 2 nM for its specific site (Ferreiro et al., 2000), which means that under these micromolar concentration conditions, E2C will stoichiometrically bind to DNA_p53_. In other words, although very low concentrations of DNA_p53_ will preferentially bind to p53, at higher concentrations DNA_p53_ will also bind to E2C. Finally, we confirmed that DNA_p53_ is present in the aggregates formed at 1:1:4 DNA:p53:E2C ratio, as judged by incorporation of FITC-labelled DNA (Figure S6B). Kinetic observation revealed that the aggregates formed at sub-stoichiometric concentration of DNA_p53_ increased in size and exhibited a wider coverage of the plate bottom concomitantly with time elapse (Figure S6C), analogous to the size increase of the heterotypic droplets without DNA.

We next wanted to evaluate the effect of long stretches of DNA that better approximates the DNA encountered by the proteins within the nucleus. To this end, we made use of calf thymus DNA (ctDNA), consisting of fragments of an average 1000 base pairs, and we represented the stoichiometry as number of 20 base pair non-specific sites along the entire fragment, approximately 50 sites. Sub-stoichiometric amounts of ctDNA deformed the homotypic p53 droplets and gradually led to their dissolution (Figure 8A). A residual of p53 droplets observed may be due to the fact that the molar ratio of DNA is not high enough but can also represent coexisting species of droplets that are more rigid and not affected by non-specific DNA, or that the inner dense phase is not accessible to a 1000 bp DNA. The effect of ctDNA was also evaluated on the heterotypic condensates, where the gradual increase of the DNA concentration led to condensation-coalescence of the initial E2C:p53 droplets into much larger irregular droplets containing both proteins, which ultimately evolved into larger condensed albeit irregular aggregates (Figure 8B).

**Figure 8.**
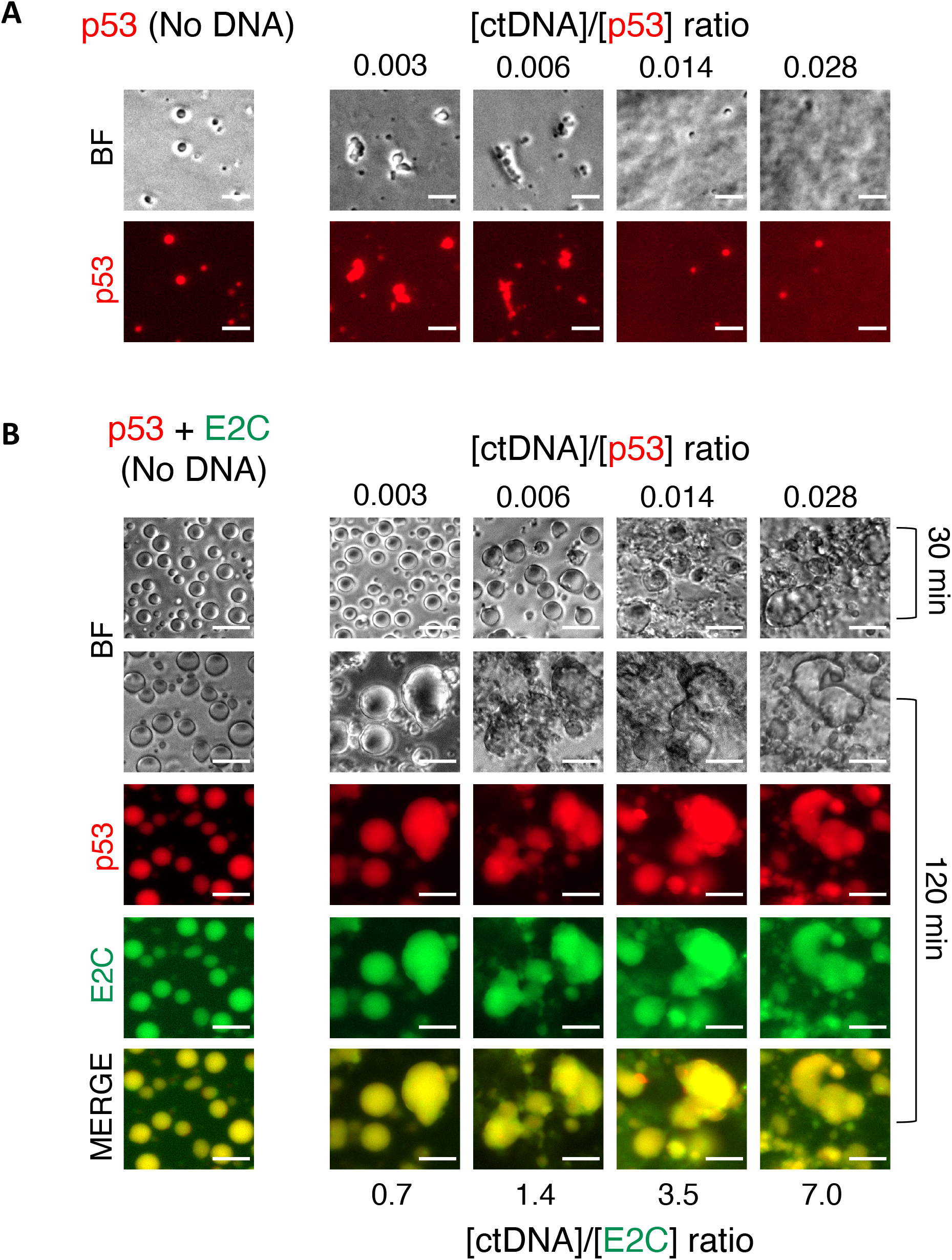
Effect of long DNA on LLPS. (A) Representative microscopy images of samples containing pre-formed p53 droplets followed by addition of increasing concentrations of calf thymus DNA (ctDNA). Control sample without DNA contains 2.5 μM cy5-p53. (B) Microscopy images of samples containing pre-formed p53:E2C heterotypic droplets followed by addition of increasing concentrations of ctDNA. Control sample without DNA contains 2.5 μM cy5-p53 and 10 μM FITC-E2C. Comparison of BF images at 30 min and 60 min, depicts the effect of time on the stoichiometry of DNA:protein complexes. Yellow regions show colocalization of p53 and E2C in the merged images. Scale bars= 10 μm.

## DISCUSSION

In this work we showed that a pseudo wild-type variant of full-length p53 can undergo homotypic and heterotypic LLPS, with functional implications for the role of p53 as inhibitor of virus replication and in connection to its loss of tumor suppression activity related to its aggregation tendencies in cancer development and progression. Homotypic LLPS of p53 yields small droplets of liquid nature in the absence of crowding agent but large and regular droplets in the presence of PEG. Both processes are reverted by salt, indicating the contribution of ionic interactions, as it was observed in a multimutated p53 (Kamagata et al., 2020). Overall, the crowder enhanced droplets reached much larger sizes, and were more spherically regular and less densely labelled, suggesting different material properties (Figures 1B, 1C and 2B). We also demonstrated the functional heterotypic condensation of p53 with HPV E2 through its DNA binding C-terminal domain (E2C). This process leads to the formation of highly regular spherical droplets that increase in size with increased concentrations of E2C, with complete colocalization of both proteins. The onset of droplet formation is at a 2:1 E2C:p53 ratio, it takes place in the absence of added crowder, and is inhibited by ionic strength, suggesting similar electrostatic components to that of homotypic LLPS, namely, interaction between highly positively charged DNA binding domains of both proteins with the negatively charged p53_TAD_. Likewise, FRAP experiments indicate that p53 displays the same dynamic properties both in homotypic and heterotypic LLPS, with a large percent of recovery. Interestingly, within the heterotypic droplets at a 4:1 E2C:p53 ratio, recovery of E2C was much faster than that of p53 displaying *t*_1/2_ = 31 seconds and *t*_1/2_ = 62.8 seconds, respectively. This finding supports the participation of E2C as client protein and, although it cannot undergo LLPS by itself, it may form weak self-interactions and diffuse more freely within the dense phase (Banani et al., 2017). Sub-stoichiometric recruitment of E2C to the PEG homotypic droplets (Figure 4C), together with its inability to form condensates on its own, also supports a role as a client. However, the effect on droplet size increase with E2C concentration is indicative of condensate modulatory activity (Banani et al., 2016; Ferrolino et al., 2018; Ruff et al., 2021), ascribing a role as an unorthodox client with strong modularity activity possibly provided by its dimeric nature and DNA binding and therefore ionic interaction capacities. However, even if the E2C domain alone can produce this effect in full, E2 has an independent transactivation N-terminal domain, separated by a disordered linker, that binds the replication essential E1 DNA helicase, and needs to be investigated in its full-length version.

We found that the spontaneous formation of the homotypic p53 droplets upon transfer to room temperature is followed by a slow gradual formation of bead-like aggregates, which appear to form by incomplete coalescence. The presence of crowder prevents the formation of these aggregates, even after 24 hours (Not shown). The time dependent formation of these homotypic aggregates is completely prevented by the presence of stoichiometric amounts of a 2:1 E2C:p53 ratio. Moreover, the p53 aggregates are progressively reshaped and converted into highly regular droplets after addition of E2C, in what can be considered a time and E2C concentration dependent rescuing mode (Figure 5A). While heterotypic p53:E2C droplets are formed in less than a minute after mixing both components (Figure 4B), reshaping of 2-hour pre-incubated p53 homotypic droplets into regular spherical heterotypic droplets by addition of E2C takes an hour to complete (Figure 5A). The source of this energy barrier appears to be in the rate of incorporation of E2C to the dense phase of an otherwise matured homotypic p53 droplet (Figure 5A). Further, when p53 is added to a solution of E2C, the homotypic p53 droplets are formed immediately and E2C is fully incorporated over a period of an hour. In other words, even when both proteins are present after initial mixing, homotypic p53 droplets are favoured kinetically against the heterotypic droplets.

The homotypic p53 aggregates strongly bound ThT, in agreement with previously described amyloid routes for aggregation of the p53_DBD_ (Ishimaru et al., 2003). Interestingly, although heterotypic droplets remained highly regular and spherical after four hours of incubation, p53 aggregation led to discrete bodies partitioned within the droplets. While an intense p53 (Cy5) and ThT labelling can be observed in these aggregates, there is also a diffuse staining of both dyes in the rest of the droplet. This observation supports the presence of two populations of conformers both capable of binding ThT, one more soluble and diffusible within the dense phase and the other a compact amyloid aggregate with more intense fluorescence labelling. We speculate that these correspond to intermediates on the amyloid aggregation route (Sivanesam and Andersen, 2019) that are first condensed and distributed homogeneously within the droplet and subsequently evolve and partition to compact amyloid aggregates (Lin et al., 2015; Wegmann et al., 2018), favoured by protein concentration and time.

Both p53 and E2 are transcription factors that operate within the nucleus where DNA is omnipresent. In this confined environment, tight_-_specific as well as non-specific DNA binding is expected. A minimal 20 bp DNA duplex with a specific p53 binding site reshaped the homotypic condensates at sub-stoichiometric concentrations and subsequently dissolve them to completeness at a 1:1 DNA_p53_:p53 ratio. This is consistent with the role of the positively charged p53_DBD_ in driving homotypic condensation through ionic, intermolecular interaction with the p53_TAD_. A similar mechanism was described for NPM1 protein, an essential oligomeric scaffolder for nucleoli LLPS (Mitrea et al., 2018). Stoichiometric reshaping and dissolution is also observed for heterotypic p53:E2C by the DNA_p53_ duplex which is expected to bind both p53 and E2C at the micromolar concentrations used in the experiments (Ferreiro et al., 2000). Based on the sequence specificity at low nanomolar concentrations, we expect that sub-stoichiometric DNA_p53_ is preferentially bound to p53, without affecting its interaction with E2C in either aggregates or heterotypic droplets. Excess of DNA_p53_ most likely binds to E2C (*K*_D_ 0.2 nM), disrupting its interaction with p53_TAD_ not likely to have such a high affinity, causing dissolution of the droplets. Indeed, we have described that E2 can bind to non-specific DNA at low micromolar concentrations, the range used in these experiments (Ferreiro et al., 2000).

Both homo and heterotypic condensates are suppressed by salt, and E2C is highly positively charged at its DNA binding site, which suggests that its participation in the condensate network is to a great extent through interaction with the negatively charged p53_TAD_, an interaction that would be disrupted by DNA binding. The fact that both condensates are dissolved by DNA through initial deformation of the droplets as an intermediate stage is intriguing and we hypothesize that the reshaping involves DNA diffusion into the condensate and binding, and a subsequent slow gradual change in the material properties of both types of condensates before complete dissolution (Model scheme in figure 9A). A similar situation is observed with the long ctDNA fragment of 1kb, but the reshaping does not lead to dissolution but to the formation of amorphous aggregates in the case of p53 alone, and irregularly shaped condensates resulting from the coalescence of the heterotypic droplets (Model scheme in figure 9B). These heterotypic condensates are definitely not amorphous aggregates or precipitates and are compatible with the condensed material observed in cells in the chromatin fraction. Moreover, in a cell line harboring a p53 mutation known to be impaired for DNA binding, p53 localizes to the nucleus but does not remain in the chromatin fraction unless E2 is present. This indicates that this mutant p53 cannot bind strongly to the chromatin through its DNA binding site but does so through an interaction with E2, as observed *in vitro*.

**Figure 9.**
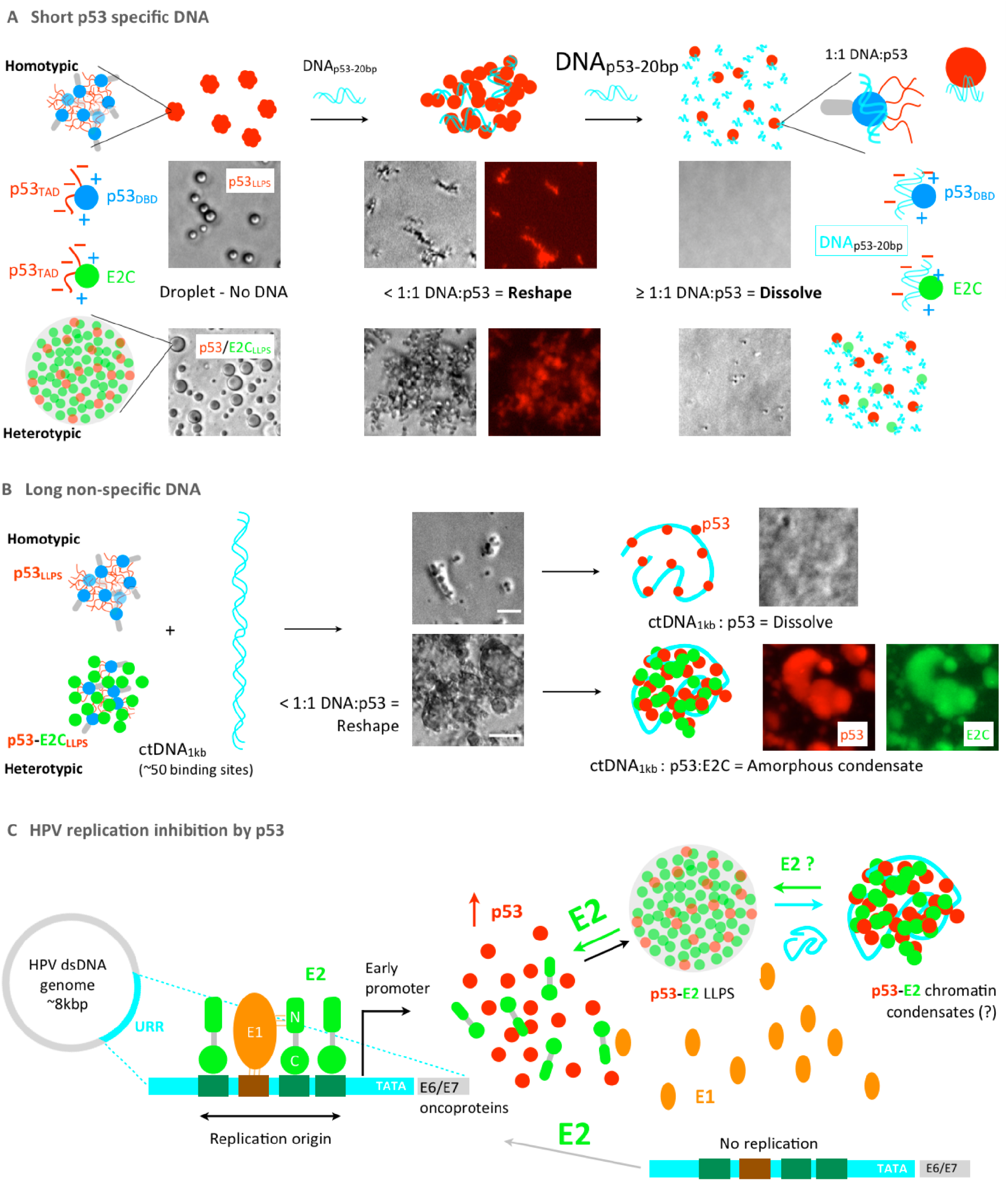
Model scheme for the effect of DNA on homotypic and heterotypic condensation and aggregation of p53 and E2C, rescuing mechanisms of p53 by HPVE2C, and HPV genome replication inhibition by p53. A) p53 20 bp binding site duplex (DNA_p53-20bp_). The DNA_p53-20bp_ oligo diffuses into the droplets and reshape it to aggregates at sub-stoichiometric ratios. At over-stoichiometric ratios (>1:1 p53:DNA) the droplet is dissolved and all p53 is bound to DNA_p53-20bp_. The DNA_p53-20bp_ also binds E2 and competes with the negatively charged p53_TAD_ for the DNA binding sites both proteins. B) Effect of the ∼1kb calf-thymus DNA fragment (ctDNA_1kb_). ctDNA_1kb_ reshapes the homotypic p53 droplets at sub-stoichiometric ratios, assuming ∼50 20pb long non-specific sites along the 1 kb fragment. In excess of sites, the droplets are dissolved (top branch). Heterotypic droplets are reshaped into aggregate-like structures, but further addition of ctDNA_1kb_ brings about the formation of anomalous irregular condensates, which are neither droplets nor amorphous aggregates. C) HPV genome replication origin requires the E1 helicase which is recruited to its binding site by E2 bound to adjacent high affinity DNA binding sites, located within the upstream regulatory region (URR). Excess of p53 – product of HPV infection-induced stress – interferes with E2 binding to E1 and DNA, sequestering E2 into condensates with consequent impairment in viral replication and transcription. As E2 levels increase the process may potentially be reverted and gene function restored. Top, blue circles correspond to p53 DBD, red chains to p53 TAD, grey cylinders to p53 tet domain, and green circles to E2C. Bottom, p53 is represented as red circles and short and long DNAs in turquoise.

Biomolecular condensation is not only revolutionizing our understanding of intracellular compartmentalization of biochemical reactions in cell physiology and pathology, but this mechanism extends to the viral world, in particular to replication and transcription (Etibor et al., 2021; Lopez et al., 2021; Nevers et al., 2020). Emerging common strategies to eukaryotic gene function involve the formation of transcription factories through the assembly of phase-separated condensates as super-enhancers or repressors (Peng et al., 2020; Safari et al., 2019). Likewise, evidence of DNA viral replication compartments of liquid-like nature (Hidalgo et al., 2021; Seyffert et al., 2021) might impact the virus-host interaction landscape. Replication was shown to take place through the formation of viral replication centers (VRCs) or foci containing the required components in different double stranded DNA viruses (Charman and Weitzman, 2020), and these were described in detail for papillomaviruses (Khurana et al., 2021; Sakakibara et al., 2011). Intriguingly, the Epstein-Barr nuclear antigen 1 (EBNA1) and latency associated nuclear antigen (LANA) of Epstein-Barr virus (EBV) and Kaposi’s sarcoma herpes virus (KSHV) respectively, not only share functional features with HPV E2, but share a unique folding topology (De Leo et al., 2020). Interestingly, LANA nuclear bodies (LANA-NBs) which are highly stable during latency showed partial dynamic structural properties involving phase separation condensates (Vladimirova et al., 2021). Moreover, LANA was shown to interact and trigger degradation of p53 (Cai et al., 2006). We hypothesize that a plausible mechanism for p53-mediated inhibition of HPV replication and transcription takes place through sequestration of E2 into highly stable heterotypic condensates that we observed in cells (Model summarized in figure 9C). Although p53 can inhibit HPV replication, recruitment of p53 to replication foci could be beneficial for the virus. E1/E2 foci represent viral replication factories and recruit DDR proteins including phosphorylated p53 at serine 15 (Khurana et al., 2021; Sakakibara et al., 2011). Also, upon early infection the capsid protein L2 interacts with E2 and recruits it to PML-NBs, and this could be important for initiation of viral replication and transcription (Day et al., 1998). Recently, it has been shown that the DDR component 53BP1 forms phase-separated compartments that are enriched in p53 (Kilic et al., 2019). In this context, the recruitment of p53 and maybe other DDR factors by E2 in the LLPS scenario may serve to repair the viral DNA or to achieve accurate viral replication.

Given p53’s features, its strong tendency to undergo LLPS is not surprising. These include, modularity, oligomerization, nucleic acid binding, IDRs, charge distribution and regulation by posttranslational modifications (Krois et al., 2016; Sun et al., 2021). This provides a conceptual understanding to its role as cell signaling hub, capable of binding a myriad of protein ligands (Hernandez Borrero and El-Deiry, 2021). We have shown that homotypic p53 LLPS is an intermediate state on the pathway to aggregation, linking LLPS to the increased tendency of frequent hot-spot p53 LOF cancer mutations to follow this route. The fact that E2 not only prevents aggregation but also rescues p53 from aggregation provides an additional angle to unveil mechanisms that can be targeted by anticancer p53 stabilizing drugs.

## MATERIALS AND METHODS

### Expression and purification of recombinant proteins

The HPV-16 E2 C-terminal domain (E2C) was expressed in *Escherichia coli* BL21(DE3) and purified as described previously (Ferreiro et al., 2000; Smal et al., 2009). The pET24a plasmid containing the sequence of stable mutant of full-length p53 with the following mutation in the DBD: M133L/V203A/N239Y/N268D (Nikolova et al., 1998) was a kind gift from Alan Fersht. These mutations conserve the functional and conformational properties of wild-type p53, allowing increased levels of expression and sample stability. Therefore, it is considered as pseudo wild-type p53 protein and will be referred as p53 throughout the text. p53 was expressed in *Escherichia coli* C41 and purified by using standard His-tag purification protocols, followed by tobacco etch virus (TEV) protease digestion and heparin affinity chromatography. The final purification step was size exclusion chromatography (SEC) using Superdex-200 column (Cytiva). p53 tetrameric conformation was confirmed by estimating the hydrodynamic radius by dynamic light scattering (DLS). Protein concentration was determined by UV light absorbance at 280 nm using the protein molar extinction coefficients and confirmed by Bradford assay. Purified proteins were stored as at −80 °C after snap freezing in liquid nitrogen and aliquoted in fixed volumes to avoid repetitive thaw-freezing. Far-UV circular dichroism and fluorescence spectra, and SDS-PAGE were performed to confirm the quality and purity of the proteins, respectively.

### Fluorescent labeling

p53 and E2C proteins were labeled with fluorescein isothiocyanate (FITC) (Sigma-Aldrich), adapting the manufacturer’s protocol to obtain sub-stochiometric labeling enough to visualize the samples by fluorescence and confocal microscopy. A p53/FITC ratio of 2 and a E2C/FITC ratio of 6 were used. Reactions were carried out at 4 ºC overnight in potassium phosphate buffer 50 mM, NaCl 0.3 M and 1 mM DTT pH 7.0. The reactions were stopped using 50 mM Tris-HCl pH 8.0 and excess of FITC removed by desalting PD10 columns (G&E) eluting each protein with the corresponding stock buffers. A similar procedure was carried out for labeling p53 with cy5 NHS (Lumiprobe) using a p53/cy5 ratio of 2. These protocols yield proteins stocks labelled with 10% - 20% fluorescent dye.

### DNAs

Double-stranded 26 bp oligonucleotides containing p53 consensus sequence (el-Deiry et al., 1992) were prepared as follows: single-stranded oligonucleotides were purchased, HPLC purified, from Integrated DNA Technologies (Coralville, IA). p53_DNA_A: 5’ AGC TT *AGGCATGTCT AGGCATGTCT* A 3’ and p53_DNA_B: 5’ AGC TT *AGACATGCCT AGACATGCCT* A 3’ (Recognition sequences are italicized). Single-stranded oligonucleotide concentration was calculated using the molar extinction coefficient obtained from the nucleotide composition. Annealing was performed as described previously (Ferreiro et al., 2000). This yielded double-stranded oligonucleotide termed DNA_p53_, and no detectable single-stranded oligonucleotide was judged by PAGE. Calf thymus DNA (ctDNA) (Sigma-Aldrich) was dissolved following the manufacture’s instructions to obtain a 1 mg/ml stock solution. ctDNA quantification was confirmed by nanodrop (Thermo Scientific) and quality was assessed by agarose gel electrophoresis.

### Light scattering kinetic measurements

Complexes formed by E2C:p53 condensation were recorded in a Jasco UV spectrophotometer by following scattering signals at 370 nm. Measurements were performed in 50 mM Tris-HCl (pH 7.0), 40 mM - 200 mM NaCl (depending on the condition) and 1 mM DTT. When crowding was needed, 5-10% PEG4000 was used. Measurements were conducted in different tubes containing the same p53 concentration (between 0.25 μM and 1 μM) and varying E2C concentrations. The 280 nm absorbance of soluble fractions was measured after centrifugation of the samples. This was done in the same apparatus. Then the pellet fraction and soluble fraction were precipitated with TCA and analyzed by SDS−PAGE.

### Turbidity assays

Protein samples were prepared in 50 mM Tris-HCl buffer, 40 mM NaCl, and 1 mM DTT pH 7.0 or 50 mM Tris-HCl buffer, 150 mM NaCl, 1 mM DTT, and 10% PEG, pH 7.0, in a total volume of 100 μl. The samples were incubated at room temperature for 15 min, and absorbance at 370 nm was measured using a Beckman Coulter DTX 880 multimode detector. All the conditions were measured in 96-well plates (Corning nonbinding surface). Each turbidity assay was reproduced in triplicate.

### Bright field microscopy and fluorescence microscopy

Analyses of homotypic p53 LLPS and heterotypic p53:E2C LLPS were performed using bright field and florescence microscopy, under varying conditions, including concentration, ionic strength, and molecular crowding. Samples were prepared at room temperature using 0.3 μM labeled protein within the bulk of unlabeled materials and loaded into 96-well plates (Corning nonbinding surface). Sample buffer consisted of 50 mM Tris-HCl buffer, 40 mM and 1 mM DTT pH 7.0. When molecular crowding conditions were needed, sample buffer consisted in 50mM Tris-HCl buffer, 150 mM NaCl, 1 mM DTT and 10% PEG pH7.0. For fluorescence analysis, the samples were excited at 488 nm for FITC or 633 nm for cy5. The images were acquired using an Axio Observer 3 inverted microscope with a 40x/0,750.3 M27 objective and a Colibri 5 LED illumination system. Images were processed using Fiji (A distribution package of ImageJ software, USA).

### Fluorescence recovery after photobleaching

Samples were prepared and imaged using Nunc Lab-Tek Chambered Coverglass (ThermoFisher Scientific Inc) at room temperature, with 0.3 μM labeled samples within the total of protein concentration. Fluorescence recovery after photobleaching (FRAP) experiments were performed using a Zeiss LSM 880 Airyscan confocal laser scanning microscope with a Plan-Apochromat 63/1.4 objective lens. For homotypic and heterotypic droplets a circular region of interest (ROI) (diameter ∼ 1 μm) was bleached using 90% laser power. Droplets of an approximate diameter of 2 μm and 4.5 μm were selected for homotypic and heterotypic condensates, respectively (7-8 n). Fluorescence intensity changes with time were recorded for three different ROIs (bleached droplet, reference droplet, and background) over 300 frames (300-350 sec), including 10 frames before bleaching. Acquired images were 512 × 512 pixels, with 1.35 μsec pixel dwell and 120.48 msec scan time. The fluorescence intensities from bleached ROI were corrected for photofading and normalized with respect to the bleaching depth as described (Alshareedah et al., 2021). Fluorescence recovery data were evaluated using the FIJI ImageJ software. Data was fit to the single exponential recovery function *y* = *A* x exp(−*k x t*) using profit 7.0.18 Quantum Soft. To improve the goodness of the fit, two-exponential fit *y* = *A1* x exp(−*k1 x t*) + *A2 x exp*(−*k2 x t*)) was also used, where *A* and *k* are fitting parameters. Half time of recovery (*t*1/2) was obtained graphically for the latter.

### Cell culture and transient transfection

All cell lines were obtained from the ATCC. C-33A cells were maintained in Dulbecco’s Modified Eagle’s Medium (DMEM) (SIGMA) supplemented with 10% FBS and penicillin-streptomycin (100 units/ml and 100 μg/ml respectively) at 37°C in 5% CO_2_. Saos-2 cells were maintained as above but in the presence of 2 mM L-glutamine. C-33A cells were transiently transfected using FuGENE 6 (Roche) following the manufacturer’s protocol. Saos-2 cells were transiently transfected using GeneJuice (Merck/Novagen).

### Antibodies

The following antibodies were used: rabbit polyclonal Lamin A/C (H-110 Santa Cruz), mouse monoclonal α-Tubulin (Santa Cruz), rabbit polyclonal p53 (FL-393 Santa Cruz). Goat anti-rabbit IgG-TRITC (Sigma-Aldrich) was used as secondary antibody.

### *In situ* subcellular fractionation

Cells growing on coverslips were washed twice with 1X PBS before cytoplasmic and loosely held nuclear proteins were removed using 200 μl of CSK I buffer (10 mM PIPES, pH 8.0, 300 mM sucrose, 100 mM NaCl, 3 mM MgCl_2_, 1 mM EGTA, pH 8.00 containing 1% (v/v) Triton X-100) for 1 min at 4°C. The coverslips were washed twice with ice cold 1X PBS and the chromatin fixed using 4% formaldehyde for 20 min at 4°C. α-tubulin and lamin A/C were visualized to confirm protein extraction.

### Immunofluorescence

Cells growing on coverslips were washed twice with 1X PBS and fixed in 4% formaldehyde at 20°C for 30 min followed by 3 washes with 1X PBS. After fixation, cells were permeabilized by incubation with 0.2% Triton X-100 on ice for 15 min and washed twice with 1X PBS. To reduce background staining the cells were incubated with 3% BSA at 20°C for 20 min and washed twice with 1X PBS. The cells were incubated with primary antibody at 4°C for 1 hour and washed twice with 1X PBS and then incubated in the dark with secondary antibody at 22°C for 1 hour and washed twice with 1X PBS. Coverslips were mounted onto glass slides using VectaShield Mounting Medium with DAPI (Vector) and visualized by fluorescence or confocal microscopy.

### Cell imaging

Cell imaging was performed using a Leica DM IRBE inverted epi-fluorescent microscope fitted with DAPI, FITC and TRITC filter sets and a 40X objective along with the Leica IM50 Version 1.20 software (Leica) for fluorescence microscopy. A Leica DM IRBE inverted confocal microscope with a 63X objective and a TCS-NT4 software (Leica) was used for confocal microscopy. The confocal images were obtained as slices representing three averaged scans. GFP were visualized using a 488/514 nm Ar/ArKr laser (green channel), whereas TRITC was observed with a 543/594 nm HeNe laser (red channel). DNA staining was visualized at 395/405 nm (blue channel).

## AKNOWLEDGMENTS

We thank Alan Fersht for full-length p53QM expression vector. We thank Ignacio Sánchez for helpful discussions. This work was funded by ANPCyT (PICT 2019-03295). SSB, and GPG are CONICET career investigators. MS is CONICET doctoral fellow. MF, AHR, and CP a are CONICET technical staff members.

