## Supplementary figures 1-6 for "The guardian of the genome meets a viral master gene regulator at a biomolecular condensate"

^3^ Institute of Cancer and Genomic Sciences, University of Birmingham, Birmingham, United Kingdom.

**SUPPLEMENTARY FIGURES**

Figures S1-S6

**A**

**B**

**C**

10% PEG

FITC-p53

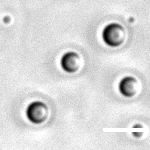

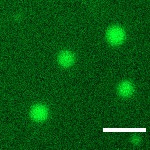

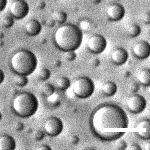

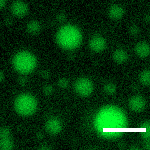

BF

p53

Buffer

p53

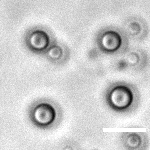

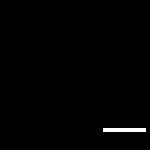

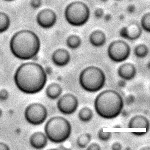

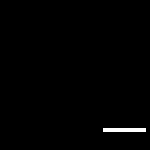

10% PEG

Buffer

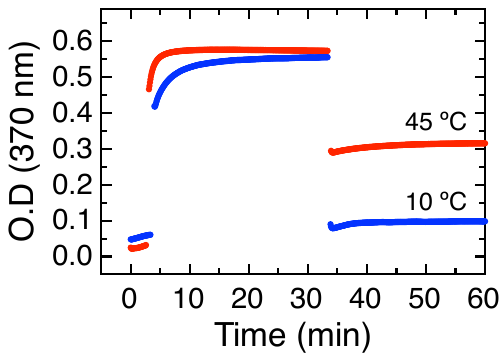

5% PEG

0.2 M NaCl

p53

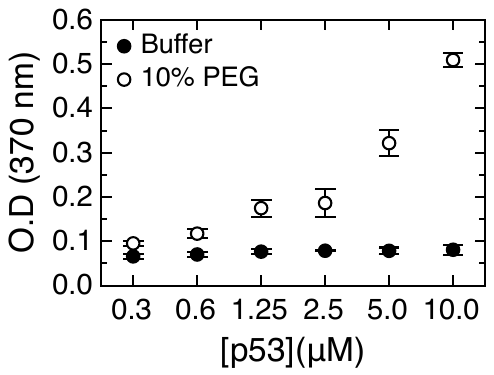

p53

**Figure S1. p53 homotypic droplets are modulated by molecular crowding and ionic strength but not affected by temperature**

(A) Turbidity kinetic assay on a sample containing 2.5 μM p53 at 10 ºC (Blue line) or 45 ºC (Red line). Addition of 5% PEG, led to a sharp increase in the absorbance signal with a more prominent effect at 45 ºC. Ionic strength considerably reduced the scattering signal, more pronouncedly at 10 ºC. (B) Concentration-dependence of p53 droplet formation was monitored by light scattering assay at 370 nm on samples with increasing amounts of p53 in absence of crowding agent or in presence of PEG. (C) Representative microscopy images of homotypic droplets formed in samples containing 2.5 μM FITC-p53 (Left panel) or 2.5 μM unlabelled p53 (Right panel) in absence or presence of PEG. BF, bright field. Scale bars= 10 μm.

**A**

**C**

**E**

**B**

**D**

p53

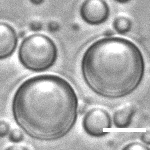

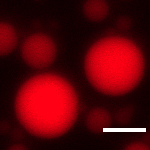

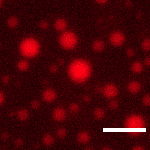

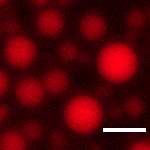

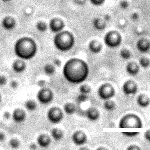

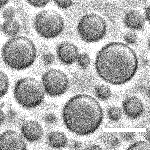

1 hour

24 hours

5 hours

10% PEG

1 hour

24 hours

5 hours

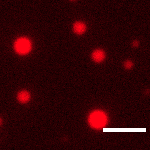

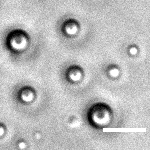

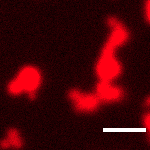

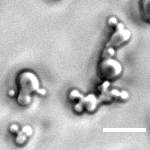

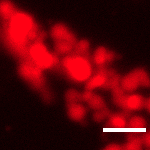

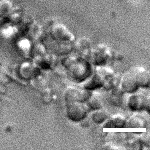

p53

BF

Buffer

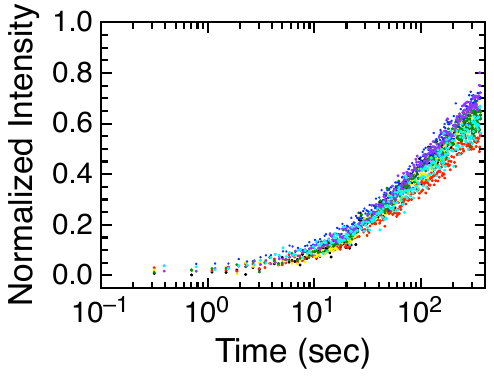

p53

p53

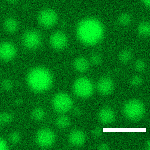

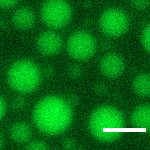

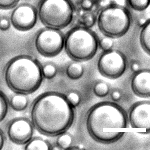

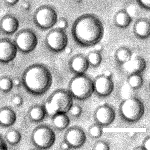

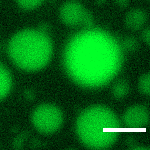

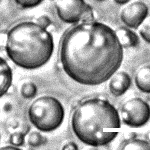

1 hour

24 hours

5 hours

10% PEG

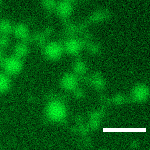

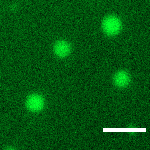

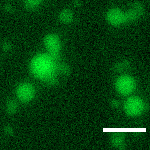

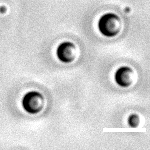

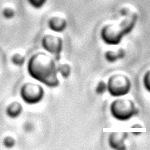

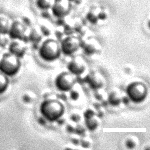

1 hour

24 hours

5 hours

p53

BF

Buffer

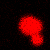

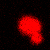

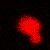

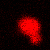

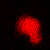

0 min

3 min

4 min

5min

6 min

10 min

p53

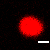

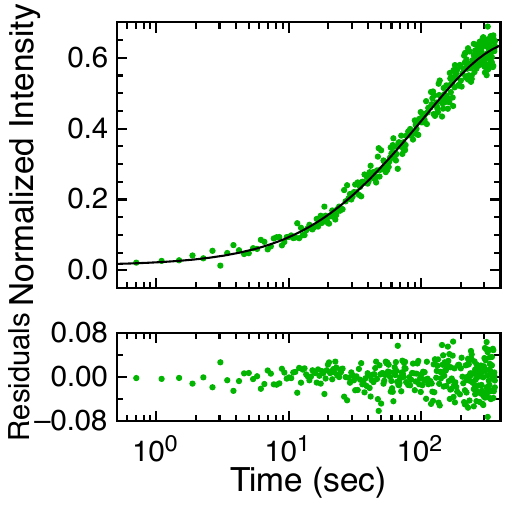

p53

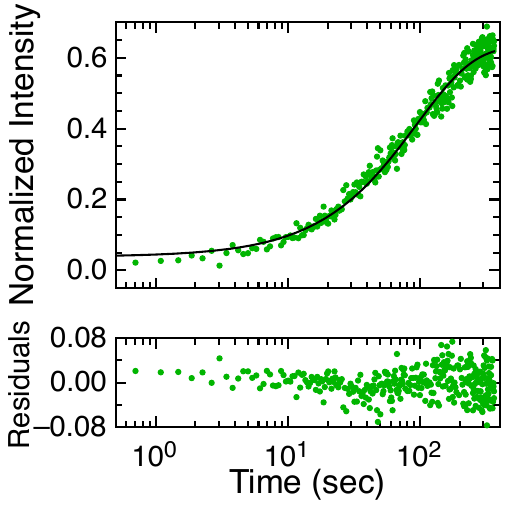

p53

**Figure S2. Fluorescence recovery after photobleaching (FRAP) analysis and time-lapse evolution of p53 homotypic droplets**

(A) Plot of FRAP analysis of FITC-labeled p53 homotypic droplets (n= 7 droplets) shows consistent replicates, indicating the reproducibility of the data. Data were normalized to the average intensity of a droplet not photobleached. (B) Normalized FRAP curves (Green dots) fitted with a single exponential function (Left plot) or double exponential function (Right plot) (Black lines). The residuals are shown in respective cases, justifying our choice of a double exponential function. A major recovery phase of 75% amplitude was obtained from the doble exponential fitting. A minor fast recovery phase was also observed but could not be explored further under the current experimental conditions. (C) Time-lapse confocal microscopy images of two homotypic cy5-labelled p53 (Cy5-p53) droplets shows incomplete coalescence leading to an irregularly shaped cluster. Scale bar= 2μm. (D) 2.5 μM cy5-p53 homotypic samples without addition of ThT (Control samples corresponding to those presented in figure 2B) showing time-course progression of spherical droplets into aggregates when crowder was not present (Left panel) or into larger regular droplets in presence of 10% PEG (Right panel). (E) p53 homotypic samples under similar conditions as in D but using FITC-labelled p53. BF, bright field. Scale bars= 10 μm.

**A**

**B**

BF

E2C

p53 + FITC-E2C

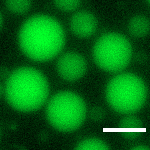

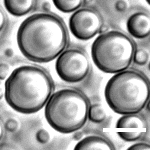

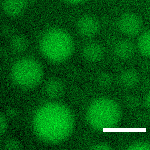

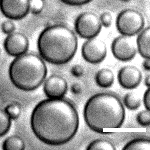

10% PEG

Buffer

FITC-E2C

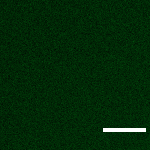

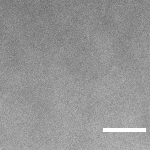

10% PEG

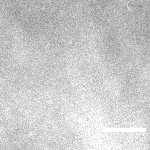

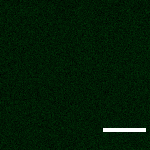

Buffer

0.25

0.5

8.0

1.0

2.0

4.0

0

[E2C]/p53] ratio

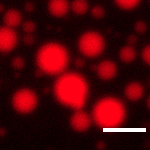

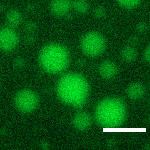

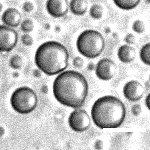

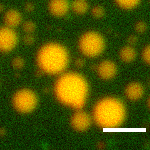

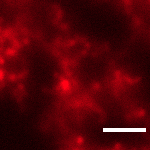

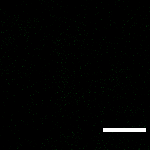

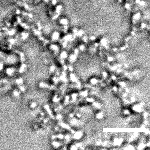

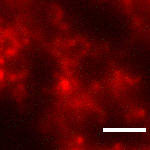

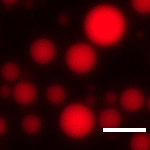

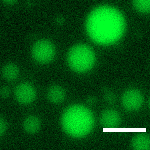

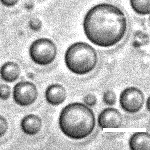

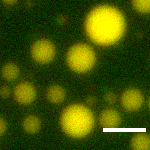

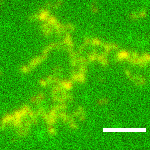

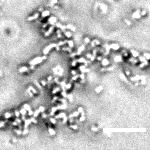

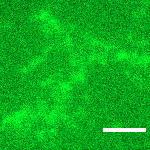

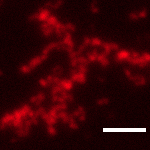

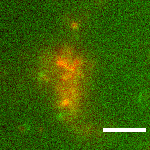

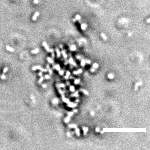

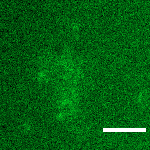

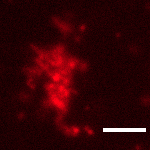

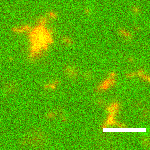

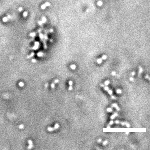

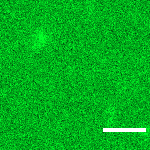

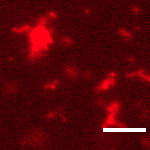

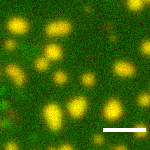

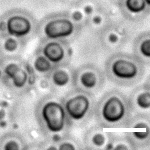

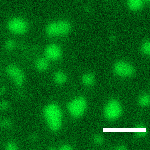

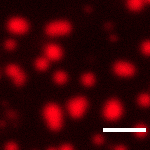

BF

p53

E2C

MERGE

p53 + E2C

0 s

1 s

2 s

5 s

p53 + E2C

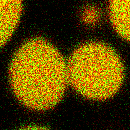

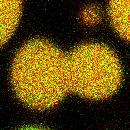

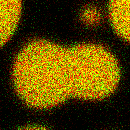

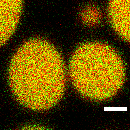

11 s

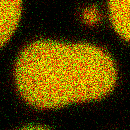

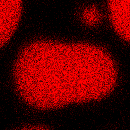

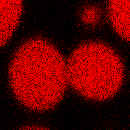

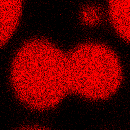

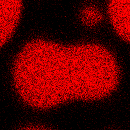

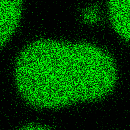

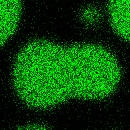

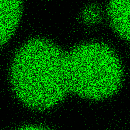

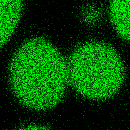

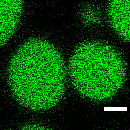

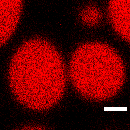

p53

E2C

MERGE

4 s

7 s

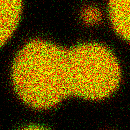

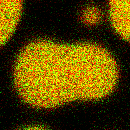

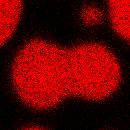

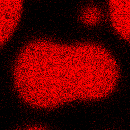

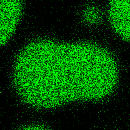

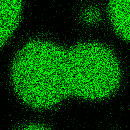

**C**

**Figure S3. p53:E2C heterotypic droplets are liquid-like and modulated by E2C availability**

(A) Combined BF and fluorescence microcopy images of 2.5 μM cy5-p53 and increasing FITC-labelled E2C (FITC-E2C) concentrations, after incubation for 7 hours. The samples are the same as those presented in figure 3C, showing that heterotypic droplets are stable at E2C/p53 ratios equal to or greater than 2.0. At ratios equal to or lower than 1.0, p53 and E2C are entangled in aggregates that gradually acquire fibrillar morphology as E2C concentration diminishes. (B) Representative microscopy images of heterotypic droplets formed in samples containing 2.5 μM unlabelled p53 and 10 μM FITC-E2C without crowding agent or with 10 % PEG (Left panel). FITC-E2C alone remains soluble even in presence of PEG (Right panel). BF, bright field. Scale bars= 10 μm. (C) Time-lapse confocal microscopy images of a typical fusion event between two heterotypic cy5-p53/FITC-E2C droplets. Scale bars= 2 μm.

**A**

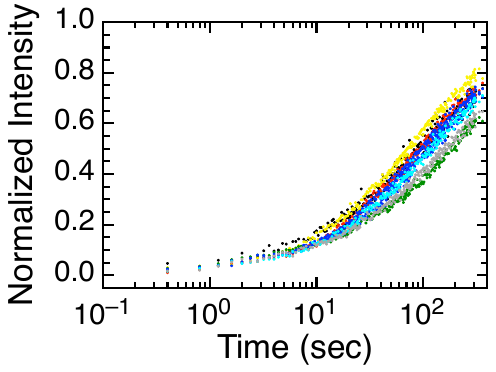

p53 + E2C

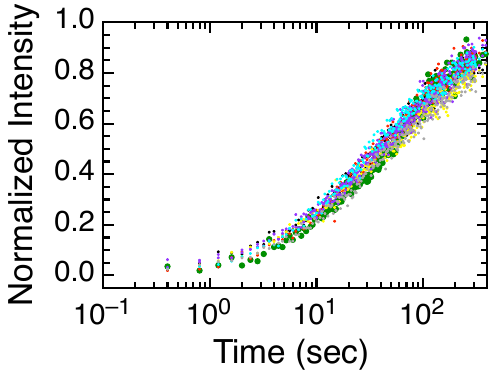

p53 + E2C

**B**

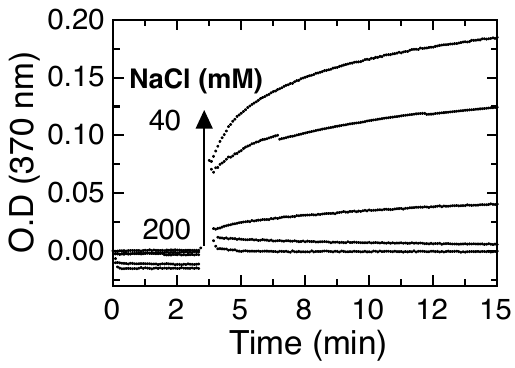

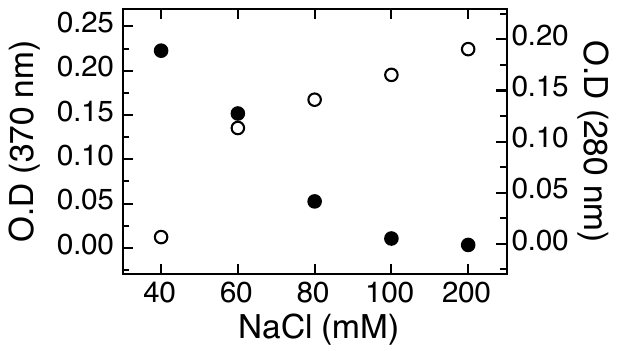

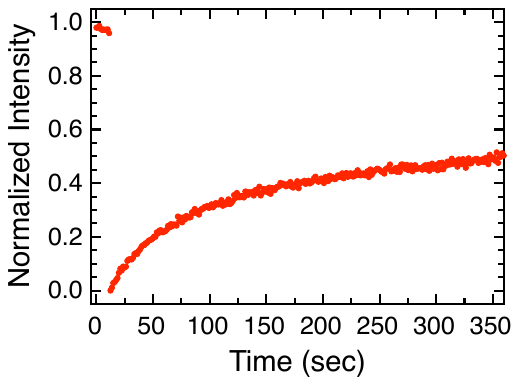

**Figure S4. FRAP behavior, coalescence and ionic strength-modulation of p53:E2C LLPS**

(A) Plot of FRAP analysis of heterotypic droplet replicates composed of 2.5 μM FITC-p53 and 10 μM unlabelled E2C (n= 7 droplets) (Left plot) or 2.5 μM unlabelled p53 and 10 μM FITC-E2C (n= 8 droplets) (Right plot). Data were normalized to the average intensity of a droplet not photobleached. (B) Kinetic light scattering plot of individual samples containing p53 pre-incubated followed by addition of E2C (E2C/p53 ratio= 2.0) with NaCl concentrations ranging from 40 mM to 200 mM (Left panel). Centrifugation of the samples was followed by measurement of absorbance signal at 280 nm (Open circles), plotted in comparison to the scattering absorbance signal at 370 nm (Black circles) (Right panel).

**A**

**B**

**C**

**D**

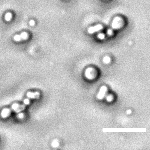

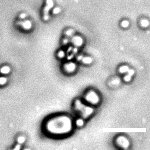

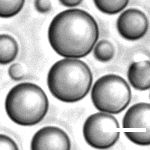

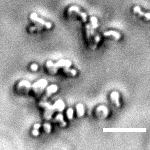

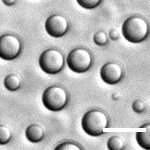

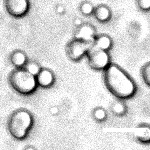

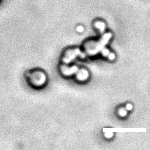

BF

20 min

40 min

60 min

20 min

40 min

60 min

Control (No E2C)

E2C

p53

(2 hour)

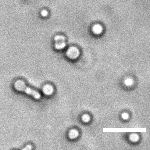

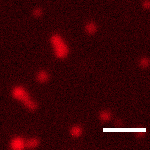

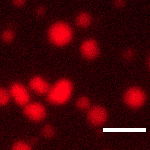

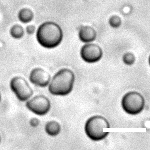

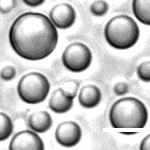

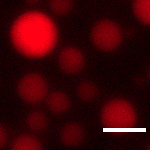

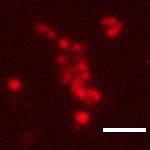

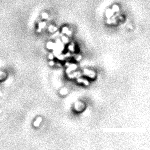

BF

p53

20 min

40 min

60 min

E2C

p53

(2 hour)

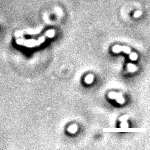

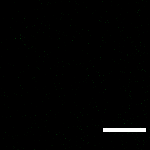

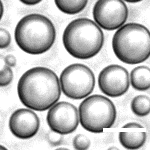

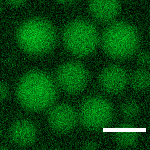

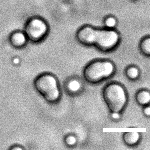

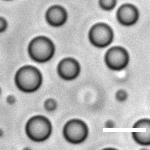

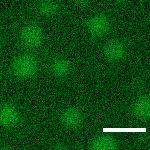

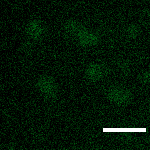

20 min

40 min

60 min

BF

p53

p53

(2 hour)

E2C

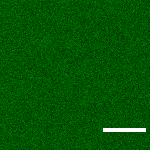

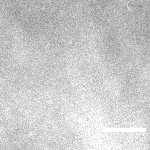

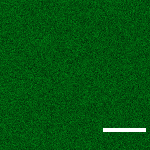

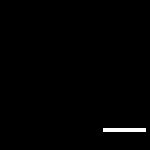

BF

E2C

E2C (2 hour)

5 min

20 min

180 min

120 min

p53

MERGE

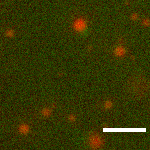

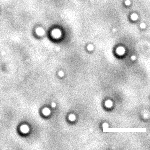

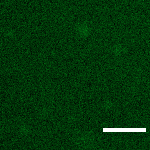

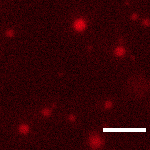

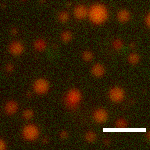

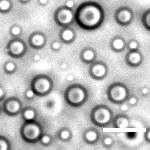

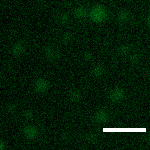

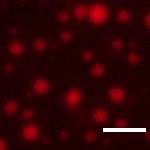

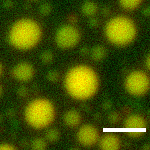

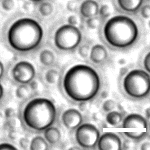

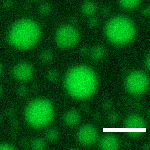

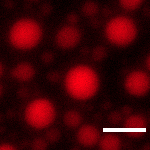

p53

**Figure S5. Evolution of p53 aggregates or E2C soluble sample into p53:E2C heterotypic droplets, upon addition of E2C or p53, respectively**

(A) Representative microscopy images of time-course examination of a sample containing 2-hour old p53 irregularly shaped clusters (Left panel) followed by addition of E2C, using unlabelled proteins. (B) Microscopy images of a sample containing 2-hour old cy5-p53 irregularly shaped clusters followed by addition of unlabelled E2C. (C) Microscopy images of a sample containing a 2-hour old p53 irregularly shaped clusters followed by addition of FITC-E2C. (D) Representative microscopy images of a sample containing 3-hour old FITC-E2C (Left panel) followed by addition of cy5-p53 (Right panel). Time-course examination shows the assemble of a phase-separated system through the formation of heterotypic droplets. BF, bright field. Scale bars= 10 μm.

**A**

**B**

**C**

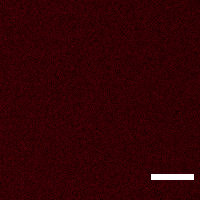

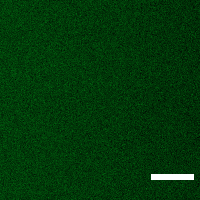

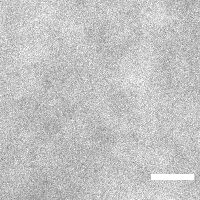

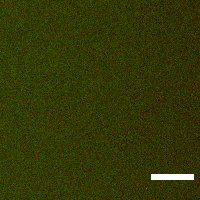

[DNA_p53_]/[p53] ratio = 1.0

p53

p53

MERGE

DNA_p53_

BF

p53

MERGE

DNA_p53_

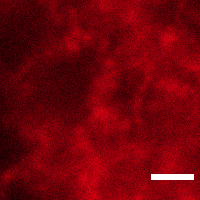

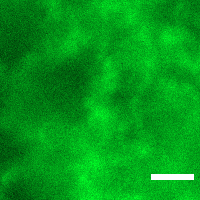

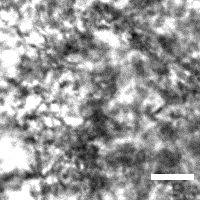

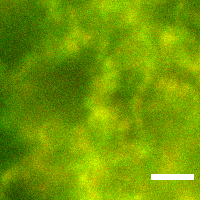

p53 + E2C

BF

[DNA_p53_]/[p53] ratio = 1.0

[DNA_p53_]/[E2C] ratio = 0.25

60 min

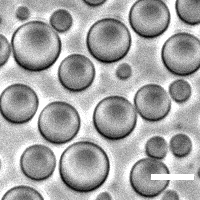

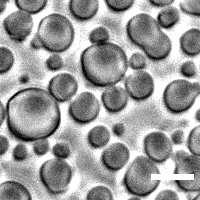

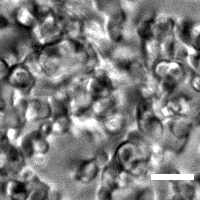

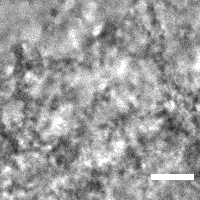

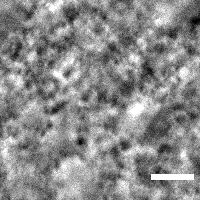

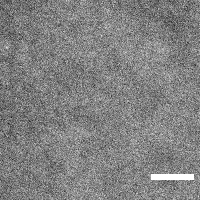

[DNA_p53_]/[p53] ratio

p53 + E2C

(No DNA)

0.4

1.0

2.0

4.0

0.2

[DNA_p53_]/[E2C] ratio

0.1

0.25

0.5

1.0

0.05

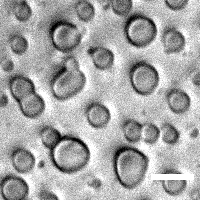

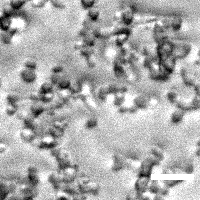

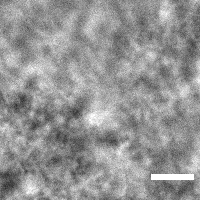

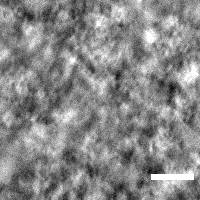

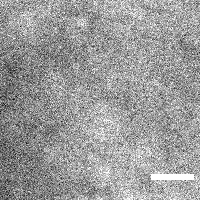

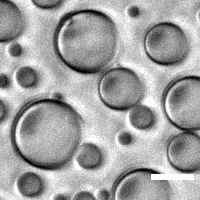

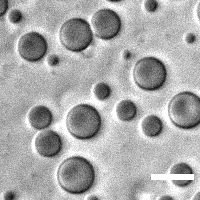

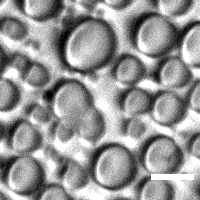

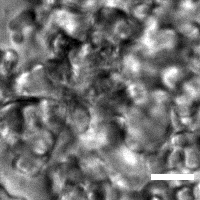

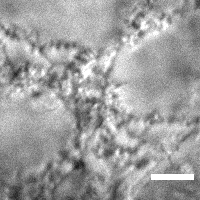

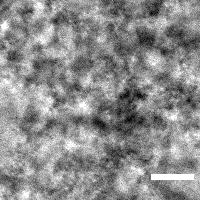

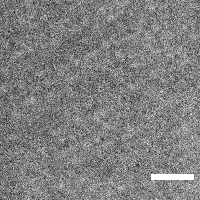

120 min

30 min

**Figure S6. p53 specific short dsDNA (DNA_p53_) is not involved in LLPS**

(A) Representative microscopy images of the effect of DNA_p53_ on p53 LLPS. Addition of 2.5 μM cy5-p53 to 2.5 μM FITC-labelled DNA_p53_ (FITC-DNA_p53_) leads to complete absence of droplets or condensates. (B) Representative microscopy images of the effect of DNA_p53_ on p53:E2C heterotypic LLPS, using 2.5 μM cy5-p53 and 2.5 μM FITC-DNA_p5_, briefly incubated before addition of 10 μM unlabelled E2C. Irregularly shaped aggregates equal to those shown on figure 7B can be appreciated, confirming the localization of both proteins and DNA_p53_ within the aggregates. Yellow regions show colocalization of p53 and DNA_p53_ in the merged images. (C) Images of time-lapse monitored samples containing 2.5 µM p53 and increasing concentrations of DNA_p53_ show that the effect of the DNA/protein stoichiometry is slightly affected by time incubation. BF, bright field. Scale bars= 10 μm.
